# Evidence that non-pathogenic microorganisms drive sea star wasting disease through boundary layer oxygen diffusion limitation

**DOI:** 10.1101/2020.07.31.231365

**Authors:** Citlalli A. Aquino, Ryan M. Besemer, Christopher M. DeRito, Jan Kocian, Ian R. Porter, Peter Raimondi, Jordan E. Rede, Lauren M. Schiebelhut, Jed P. Sparks, John P. Wares, Ian Hewson

## Abstract

Sea star wasting disease describes a condition affecting asteroids that resulted in significant Northeastern Pacific population decline following a mass mortality event in 2013. The etiology of sea star wasting is unresolved. We hypothesized that asteroid wasting is a sequela of microbial organic matter remineralization near respiratory surfaces which leads to boundary layer oxygen diffusion limitation (BLODL). Wasting lesions were induced in *Pisaster ochraceus* by enrichment with a variety of organic matter (OM) sources and by experimentally reduced oxygen conditions. Microbial assemblages inhabiting tissues and at the asteroid-water interface bore signatures of copiotroph proliferation before wasting onset, followed by the proliferation of putatively facultative and strictly anaerobic taxa. These results together illustrate that suboxic conditions at the animal-water interface may be established by heterotrophic bacterial activity in response to organic matter loading. Wasting susceptibility was significantly and positively correlated with rugosity, a key determinant of boundary layer thickness. At a semi-continuously monitored field site (Langley Harbor), wasting predictably occurred at annual peak or decline in phytoplankton biomass over 5 years, suggesting that primary production-derived OM may contribute to BLODL. Finally, wasting individuals from 2013 – 2014 bore stable isotopic signatures reflecting anaerobic processes which suggests that this phenomenon may have affected asteroids during mass mortality. The impacts of BLODL may be more pronounced under higher temperatures due to lower O_2_ solubility, in more rugose asteroid species due to restricted hydrodynamic flow, and in larger specimens due to their lower surface area to volume ratios which affects diffusive respiratory potential. Moreover, our results demonstrate that marine invertebrate disease may result from heterotrophic microbial activity that occurs adjacent to respiratory tissues, which raises important questions about the etiology of marine diseases in other benthic taxa.

## INTRODUCTION

Sea star wasting (SSW) disease describes a suite of clinical signs in asteroids including loss of turgor, arm twisting, epidermal ulceration, limb autotomy, and death. The condition gained prominence in 2013 when it caused mass mortality of >20 asteroid species in the Northeastern Pacific (Hewson et al., 2014) with continuous observations since (Miner et al., 2018;Jaffe et al., 2019). However, lesions compatible with SSW in various asteroid species have been reported since at least 1896 in the Eastern US (Mead, 1898), and at many locations globally (reviewed in Hewson et al., 2019). The cause of SSW is unresolved. Early reports that SSW was associated with a densovirus (Hewson et al., 2014) were refuted by subsequent investigation that failed to show a consistent association between the virus and presence of disease (Hewson et al., 2018), and recent description of persistent and phylogenetically widespread infection by related densoviruses (Jackson et al., 2020a;Jackson et al., 2020b) suggest this virus to be a component of normal microbiome. Furthermore, wasting is not consistently associated with any bacterial or microbial eukaryotic organism (Hewson et al., 2018). Environmental conditions, including elevated water temperatures (Eisenlord et al., 2016;Kohl et al., 2016), lower water temperatures and higher pCO_2_ (Menge et al., 2016), and meteorological conditions (Hewson et al., 2018) correspond with wasting at distinct locations. Recent modelling studies suggest repeated sea surface temperature anomalies may correlate with wasting (Aalto et al., 2020). Reports of SSW spread between adjacent geographic locations, through public aquarium intakes, and challenge experiments with tissue homogenates suggested a transmissible etiology (Hewson et al., 2014;Bucci et al., 2017). However, there is a lack of mechanistic understanding how SSW is generated in affected individuals.

Here we provide convergent evidence that asteroid wasting is a sequela of boundary layer oxygen diffusion limitation (BLODL; Fig. 1). In this model, elevated organic matter (OM) concentrations stimulate the growth of copiotrophic microorganisms adjacent to animal surfaces, driving down dissolved O_2_ concentrations and causing suboxic conditions at the animal-seawater interface within the diffusive boundary layer. Over time, these conditions do not meet respiratory O_2_ demand of tissues, resulting in their damage and decomposition, which further enriches near-asteroid pools of organic material. This in turn results in the proliferation of anaerobic taxa on and within tissues. Here we provide evidence for this effect through study of microbiome composition during organic matter amendment and as animals waste in the absence of external stimuli. Further, we demonstrate that wasting can result from suboxic water column conditions. Next, we illustrate that wasting susceptibility is related to inherent asteroid properties, including rugosity (a key physical characteristic influencing diffusive boundary layer height), animal size and respiratory demand. We explored the relationship between primary production and wasting during a 5 year time series at a field site and find significant correlation between wasting occurrence and trends in chlorophyll a, which is one potential source of OM for bacterial nutrition. Finally, we demonstrate that wasting asteroids from the 2013-2014 mass mortality event bore stable isotopic signatures that reflect anaerobic microbial processes compared to their asymptomatic sympatric counterparts.

**Fig. 1:**
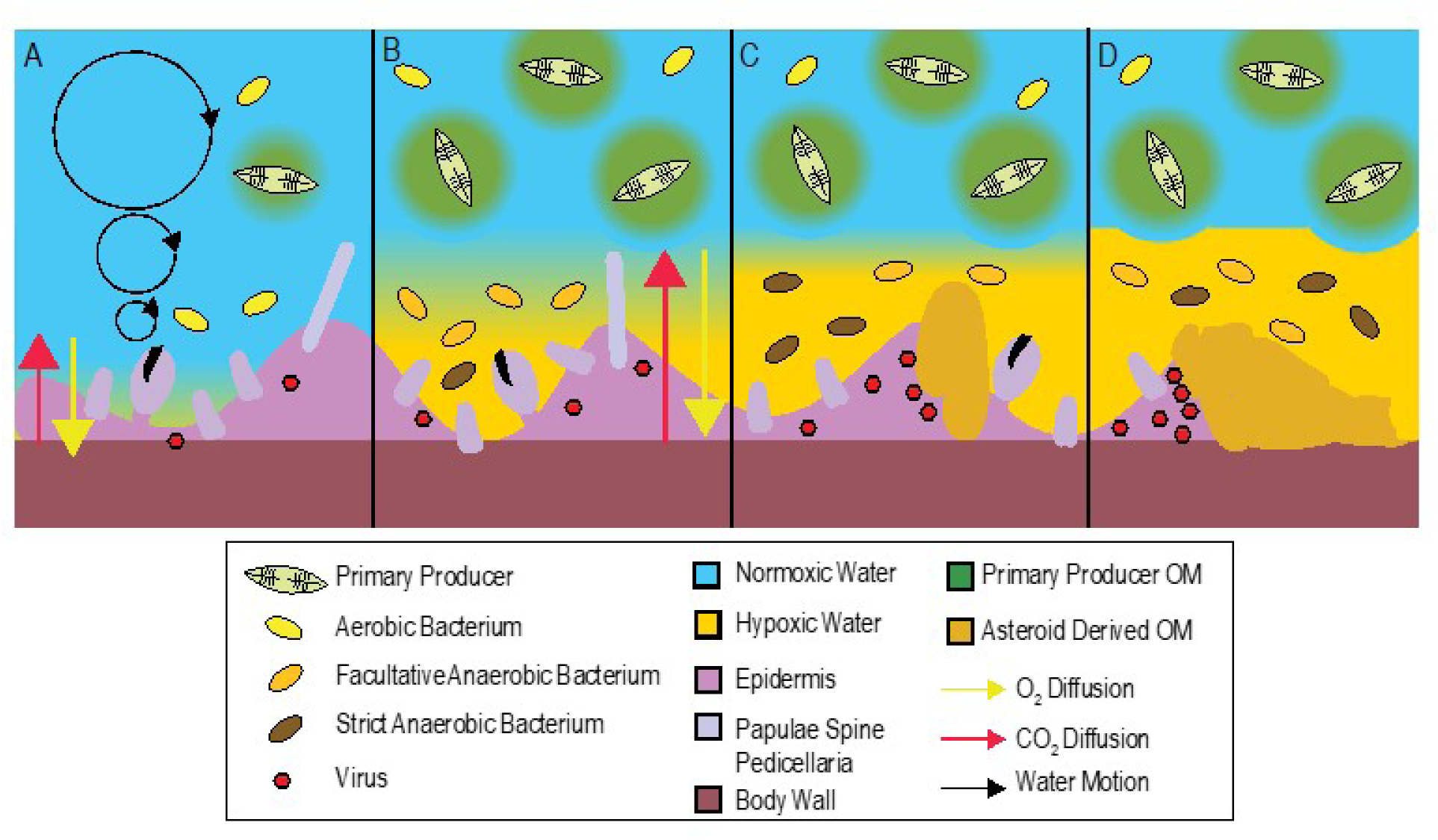
Conceptualization of BLODL. Under typical conditions (A), boundary layer conditions are normoxic because of limited dissolved organic matter inputs, and consequently lower heterotrophic bacterial respiration. When organic matter (possibly from primary production) increases in surrounding waters (B; e.g. during heightened primary productivity, from terrestrial runoff, or from decaying asteroid carcasses), this stimulates bacterial heterotrophic respiration and abundance and results in the formation of suboxic waters within the limited water motion boundary layer, which in turn results in longer distances over which diffusion must occur to maintain animal respiratory demand. Over time (C) suboxic conditions in the boundary layer results in tissue damage, and prevalence of strict and facultative anaerobes. Because their growth is less efficient than aerobic metabolisms their abundance is less than at hypoxia onset. Release of labile organic matter from decaying tissues (D) and persistent organic-matter rich conditions within the asteroid boundary layer result in animal mortality.

## METHODS

### Gross wasting definition and general specimen characteristics

Sea star wasting (SSW) in field populations is reported to comprise a wide suite of signs, including loss of turgor (deflation), discoloration, puffiness, arm twisting/curling, limb autotomy, body wall lesions and erosions and protrusion of pyloric caeca and gonads (Hewson et al., 2014). There is currently no case definition for any species of asteroid, nor is there progression of disease signs characteristic of wasting (i.e. wasting has no pathognomic signs; Hewson et al., 2019). Because many of these disease signs are subject to observer bias, we sought to standardize occurrence of wasting between experiments and surveys. We considered wasting as the appearance of non-focal body wall lesions across all experiments. We used the time between experiment initiation and body wall lesion appearance (i.e. lesion genesis) as a standardized parameter to assess the speed of wasting. Experiments were performed where individuals were physically isolated from each other while in the same aquarium water body through plastic containers which had been punched with holes (4mm diameter) at a density of ∼ 2 holes cm^-2^. However, in the May – June 2018 observation of wasting in the absence of external stimuli, individuals were identified by the unique “fingerprint” of apical circlet paxillae and spines and were not physically separated from each other. In all experiments, lesion position was standardized by clockwise naming of rays from the madreporite. Individual asteroids were considered dead when tube feet did not move during observation for 30s, or when all limbs autotomized away from the central disc. The mass of individuals was determined using a top pan balance, and each ray measured from tip to the center of central disc using a ruler or calipers.

### Impact of organic matter enrichment on wasting in Pisaster ochraceus

We examined the impact of OM enrichment on asteroid wasting to test whether enrichment caused bacterial abundance and composition shifts consistent with the boundary layer oxygen diffusion limitation (BLODL) hypothesis. Twenty *P. ochraceus* (mass 303 ± 26 g) were obtained from the jetty at Bodega Bay in July 2019 and transported to the Bodega Bay Marine Lab (UC Davis), where they were placed in flow-through large volume sea tables for 7 d prior to commencement of the experiment. Asteroids were placed individually into baskets and suspended in each of 4 sea tables with flow rates of 60 ± 15 mL s^-1^. A dense (∼10^4^ cells mL^-1^) culture of *Dunaliella tertiolecta* was prepared in 20 L artificial seawater, which was filtered onto 0.2 µm Durapore filters and resuspended in 1 L artificial seawater. The resuspended matter was divided into 21 x 45 mL aliquots and frozen prior to use. Coastal particulate OM (POM) was prepared by filtering 40 L seawater from the unfiltered intake pipe at Bodega Marine Laboratory through 0.2 µm Durapore filters, which was then resuspended in 1 L seawater. This was also divided into 21 x 35 ml aliquots and frozen before use. Individuals (n = 5 per treatment in separate sea tables) were amended with 3 OM sources: 1) peptone (approx 300 mg individual^-1^); 2) 45mL of *Dunaliella tertiolecta* filtered culture; and 3) Coastal POM (6mL) daily. Individual asteroids were observed daily (aboral and ventral surface) for the presence of lesions and they were weighed. There was no significant difference in the rate of animal mass change over the course of the experiment between treatments (Fig. S1), where individuals lost 11.0 ± 0.2 % of their initial mass over the course of the experiment. Daily samples (2 mL of water) for surface bacterial abundance were withdrawn using 3mL syringes which were pressed onto the aboral surface of individual stars while immersed, and preserved in 5% formalin and kept at 4°C in darkness prior to processing. Surface microbial communities were sampled every 48 h by collecting a swab (sterile cotton-tipped, dry transport system; Puritan) from each aboral surface which were then frozen at −20°C prior to processing. Microbiome compositional analyses was performed as described later in this section.

### Wasting progression in the absence of external stimuli

We sought to investigate longitudinal microbiome composition over wasting progression in lesion margins and in artificial scar tissues, in an attempt to discern opportunistic taxa from potential pathogens. This work was performed to target epidermis-associated microorganisms which were minimally sampled (i.e. by surface scrape), and in a separate study to target body wall tissues (i.e. by transverse body wall biopsy punch). We anecdotally observed in previous work (Hewson et al., 2018) that sampling stress, especially by collection of tissues by biopsy punch, may influence wasting in aquarium studies so sought to minimize sampling before the appearance of lesions.

To examine wasting in epidermal tissues (which presumably include microorganisms inhabiting tissue surfaces as well as those beneath the epidermis and cuticle but excluding body wall and mutable collagenous tissue-associated microorganisms), we examined temporal progression in *P. ochraceus* specimens from central California collected on 19 July 2018. Six specimens (mean mass 290 ± 54 g and ray length 11.2 ± 0.9 cm) were collected from the intertidal zone at Davenport, CA (37°1’19”N, 122°12’56”W) at low tide and transported in insulated coolers to the Long Marine Laboratory at UC Santa Cruz, where they were housed in flow-through aquaria (3.81 ± 0.05 mL s^-1^; mean water residence time 37 min) indoors in individual containers. Individuals were maintained under these conditions for the duration of the experiment. Temperature, salinity, and dissolved O_2_ were measured daily using a YSI-3000 handheld meter. After a 48 h acclimation period, a small scar (∼ 5 mm long) was made on a single ray using a sterile 4 mm biopsy punch. Individual asteroids were monitored daily for the presence of lesions (as defined above). A small tissue sample (∼ 3 mm x 2 mm) was taken from the margin of artificial scars using sterile 4 mm biopsy punches after 24 h. New artificial scars were made on adjacent rays each day, and sampled after 24h. When lesions away from these artificial scars were observed (i.e. lesions not caused by physical scarring), their margin tissues were sampled using the same approach. All tissue samples were placed into sterile 1.2 mL cryovials and immediately frozen in liquid N_2_ or in a −80°C freezer.

To examine wasting in all body wall + epidermal tissues (i.e. biopsy punch), we examined temporal progression in *P. ochraceus* specimens collected on 24 May 2018 from Davenport, CA (37°1’19”N, 122°12’56”W), which were transported in an insulated container to the Long Marine Laboratory at UC Santa Cruz where they were placed into a single, flow-through sea table. The individuals were measured, weighed and photographed to fingerprint apical circlet pattern. A single biopsy punch (4 mm) was retrieved from each individual and preserved in liquid N_2_. After 312 h, 10 of the 12 had lesions on their surfaces, and after 360 h, all individuals had lesions. After 360 h, lesions were sampled by biopsy punch and preserved in liquid N_2_, and again at time of death, which occurred between 360 h and 480 h. Because we did not capture the exact time of lesion occurrence, we restricted analysis of these individuals to samples collected at initial (T0), 360 h (TI – i.e. shortly after lesions had formed) and at time of death (TF).

### Impact of suboxic conditions on wasting in Asterias forbesi

To test whether wasting could result from suboxic conditions, we incubated asteroids in incubations where O_2_ was depleted and examined surface microbiome abundance and composition. Twenty-four *Asterias forbesi* (mass 63 ± 7 g) were obtained from Bar Harbor, ME (44°25.7’N, 68°12.0’W) on 29 June 2019 and transported in insulated coolers to the laboratory at Cornell University. There, the individuals were initially placed into a single, 320 L aquarium for 24 h, before being placed into individual flow-through baskets and divided into two treatments. One large volume (230 L) sump containing artificial seawater (Instant Ocean) was set up for each treatment. One sump served as a control, while another sump was continuously sparged with medical-grade N_2_ (∼ 5-6 L min^-1^) to lower pO_2_. Continuous flow between sump and aquarium systems was maintained by non-self-priming pumps and removed by gravity through a standpipe. O_2_ and temperature was continuously monitored in the experimental tanks using HOBO O_2_ Loggers (U26; Onset).

O_2_ was depleted in N_2_-sparged sump waters (5.87 ± 0.29 mgL^-1^) compared to control incubations (9.62 ± 0.06 mg L^-1^), representing a mean concentration decrease of ∼39%. Individual asteroids were monitored daily for the gross appearance of lesions on aboral and ventral surfaces. There was no significant difference in the rate of mass change between treatments, where all individuals lost on average 30.0 ± 6.5% of the initial mass over the course of the experiment as they were not fed (Fig. S1). At experiment initiation, surface bacterial abundance samples were taken and preserved as described above. The mass of individuals was recorded daily; stars remained without prey during the experiment. Surface microbial swabs were sampled every 48h following the approach outlined above. To reduce sampling stress on individuals, half of individuals in each treatment were biopsied (2 x 3 mm biopsy samples collected using sterile biopsy punches) on their aboral surface at experiment initiation, and then every 5 d until experiment termination (biopsied specimen lesion genesis time was not significantly different to non-biopsied specimen lesion genesis time). Biopsy punches (1 each) were preserved in RNALater or in 10% neutral buffered formalin. Upon appearance of lesions, their margins were sampled using a 5 mm biopsy punch to scrape a ∼ 3 mm x 2 mm tissue sample. Additionally, a 3 mm biopsy punch was used to obtain a sample through body wall tissues on the lesion margins. Lesion margin tissues were stored at −20°C until analysis.

### Bacterial abundance

Bacterial abundance in suboxic and organic matter amendment experiments was determined by SYBR Gold staining and epifluorescence microscopy (Porter and Feig, 1980;Noble and Fuhrman, 1998;Shibata et al., 2006). An aliquot (1 mL) of each sample was first stained with SYBR Gold (2 µl mL^-1^ of the 10,000X stock) for 2 min, then samples were filtered through 25mm diameter 0.2 µm black cyclopore filters mounted on 25 mm Type AA Millipore filters to even flow. The filters were removed from the backing filter, adhered to clean glass slides and mounted in 30 µL of PBS:Glycerol (50:50) containing 0.1% p-phenylenediamine. The slides were visualized on an Olympus BX-51 epifluorescence microscope under blue light excitation. Over 200 cells were counted in >10 fields. Bacterial abundance was calculated by multiplying mean abundance per reticle grid by total grids per filter area, and divided by volume passed through the filter. For some samples, high background fluorescence precluded accurate counts and so are not included in downstream analyses.

### Microbial assemblage analyses

Microbial assemblages inhabiting body wall samples (i.e. biopsy punch; wasting in the absence of external stimuli), lesion margins (i.e epidermal scrapes; wasting in the absence of external stimuli), and at the animal/water interface (i.e. swabs; suboxic and organic matter enrichment experiments) were examined by 16S rRNA amplicon sequencing. Nucleic acids were extracted from frozen 3mm biopsies, lesion margin scrapes, and frozen surface swabs using a Quick-DNA Fungal/Bacterial MiniPrep Kit (Zymo Research, cat# D6005) according to the manufacturer’s protocol. Bacterial DNA was quantified using a Quant-IT dsDNA Assay (Invitrogen, cat# Q33120) in conjunction with a StepOnePlus™ Real-Time PCR system (Applied Biosystems). Bacterial community composition was examined via PCR amplification sequencing of the V4 region of the 16S rRNA gene using a modified version from Caporaso et al., 2011. Fifty µl PCR reactions contained 1x 5PRIME HotMasterMix (QuantaBio, cat# 2200400) and 0.1µM each primer (515f/barcoded-806r). Template DNA quantity varied between experiments. For examining microbiome composition during wasting progression, 1 µl of extract (containing from below detection limit of 0.1 ng to 80 ng [mean = 17 ng]) was used as PCR template. For *Asterias forbesi* hypoxia and *Pisaster ochraceus* OM enrichment, 5 pg of genomic DNA (determined by Femto Bacterial DNA Quantification Kit; Zymo Research, cat# E2006) was used to standardize prokaryotic template amounts (Hewson, 2019). PCR products from duplicate reactions were pooled for each sample and cleaned using a Mag-Bind RxnPure Plus Kit (Omega Bio-tek, cat# M1386-01). Ten ng of bacterial DNA from each sample were pooled, libraries prepared using the NextFLEX prep, and sequenced on 2 lanes of Illumina MiSeq (2 x 250 paired end) at the Cornell Biotechnology Resource Center. Sequence libraries are available at QIITA under studies 12131 and 13061 and at NCBI under BioProject PRJNA637333.

Raw sequences were uploaded to QIITA and processed using the native 16S rRNA pipeline (Gonzalez et al., 2018). After demultiplexing, reads were trimmed to 150 bp and sub-OTUs (sOTUs) were configured using Deblur (Amir et al., 2017). The SEPP phylogenetic tree and BIOM/FA files were downloaded from the deblur reference hit table and converted into qiime2 (v2019.10) artifacts, after which taxonomy was assigned using the Silva 132 release (Quast et al., 2012) and q2-feature-classifier plugin. All files were then imported into R (v3.6.1) using qiime2R v(0.99) (Caporaso et al., 2010) and compiled into a phyloseq (v1.28) (McMurdie and Holmes, 2013) object for downstream analyses.

Amplicon data was transformed using the PhILR (Phylogenetic Isometric Log-Ratio Transform, v1.1) package for ordination (Silverman et al., 2017). PhILR transforms compositional data (i.e., proportional, or relative data) into a new matrix of ‘balances’ that incorporates phylogenetic information. Here, a balance is defined as the isometric log-ratio (ILR) between two clades that share a common node. The ILR is a common tool used in compositional data analysis that transforms constrained data into an unconstrained space (i.e., Euclidean space), thereby allowing standard statistical tools to be applied. For each experiment, low abundance sOTUs were filtered based on sequencing depth/evenness and a pseudocount of 0.65 was applied to all 0 counts. The ILR analyses in this study do not require an even sampling depth, and therefore, count normalization techniques like rarefying, which lead to a loss of information (McMurdie and Holmes, 2014), were not utilized. Phyloseq (1.32.0) (McMurdie and Holmes, 2013) was used to perform principal coordinate analyses (PCoA) on the Euclidean distances between PhILR transformed samples, and the adonis function in the vegan package (v2.5-6) (Oksanen et al., 2019) was used to perform a PERMANOVA on relevant PCoAs. For June 2018 *Pisaster ochraceus* samples, a principal coordinate analysis based on Weighted Unifrac distances (Lozupone et al., 2011) was used in lieu of the PhILR transformation due to higher explained variance from a PERMANOVA. For these samples, low abundant sOTUs were removed and the data was transformed to an even sampling depth. Multiple filtering strategies were applied to all analyses and did not affect results.

Balances were used to quantify differential abundance in order to address the issue of compositionality in amplicon data. Two approaches were used that rely on the ILR transformation. For comparisons between two categorical variables (diseased/non-diseased tissue and surface swabs from specimens immediately before lesions appear with earlier specimens), a sparse logistic regression with an *l*_1_ penalty of λ=0.15 was applied to PhILR balances using the glmnet package v(3.0-2) (Friedman et al., 2010). For time-course experiments, the PhyloFactor package (v0.0.1) was used. PhyloFactor calculates balances in a similar fashion to PhILR, but instead of using nodes to contrast clades, Phylofactor bisects a phylogenetic tree along its edges in an iterative manner. Each iteration, or ‘factor,’ is regressed using a generalized linear model. Edges were maximized using the F statistic and a Kolmogorov-Smirnov test was used to break the iterations.

### Additional experimental challenges of Pisaster ochraceus

We also sought further evidence of BLODL by examining wasting in context of water flow rates, desiccation stress, and challenge with tissue homogenates from a wasting specimen. *P. ochraceus* (n = 24) were collected from Mitchell Cove, Santa Cruz (36°57.1N, 122°2.51’W), on 25 June 2018 at low tide (mean mass 357 ± 37 g), and transported in a cooler to the Long Marine Laboratory. Metadata on their size and weight, along with mean flow rates in incubations and change in mass over time is provided in Table S1. The temperature in aquarium settings for all experiments was measured by Onset Hobo Spot loggers (n = 2) which were deployed into aquarium outflows for the first 4 days of the experiment, and after 6 days was measured using a YSI Handheld Instrument.

The impact of water flow on asteroid wasting was examined in 12 specimens. Asteroids (n = 6 each treatment) were placed into individual plastic boxes which were subject to high (7.06 ± 0.39 ml s^-1^; water residence time in container ∼ 20 min) and low (2.84 ± 0.26 ml s^-1^; water residence time 50 min) flow-through rates. Temperature and salinity were monitored daily using a handheld YSI Probe (YSI-3000). Individuals were visually inspected daily for the presence of body wall lesions and were weighed to determine changes in their overall mass over the course of the experiment.

The experiments were performed over a 21 day period during which mean water flow-through temperatures ranged from 14.4 – 17.6°C (Fig. S2), with day-night variation of 1.8 – 1.9°C. Water temperatures increased mostly between 10 and 15 days of incubation. Salinity did not vary by more than 0.2 over the course of the experiment. Additionally, we deployed two loggers into the intertidal zone at Davenport, CA (37°1’19”N, 122°12’56”W) to examine variation between experimental temperature conditions and conditions experienced by asteroids at the collection site. In contrast to experimental flow-through systems, field-deployed HOBO loggers revealed strong changes in temperature accompanying tidal cycles and cycles of immersion/emersion (Fig. S3). The maximum temperature variation recorded *in situ* occurred during a low tide in the morning of 29 June, when the temperature swung from from 11.0°C at 6:00am to a maximum of 33.7°C at 10:08am, then back to 13.3°C by 11:00am.

The impacts of desiccation under both high and low flow was examined by first placing 6 individuals onto plastic trays in sunlight for 1 h. During this period, air temperature was 33.8°C (mean flow-through incubation temperature was 14.5°C). After desiccation for 1 h, individuals were placed into individual flow-through plastic boxes and monitored per the variable flow rate experiments described above. Comparison between desiccation and tissue homogenate challenge (described below) were performed against controls high/low flow as described above.

The effects of challenge with wasting tissue homogenates was examined in 4 individuals. A single wasting star was collected at Davenport, CA (37°1’19”N, 122°12’56”W) at low tide on 25 June 2018 and transported to the Long Marine Laboratory. Tissue surrounding lesions (∼ 2 g total) was excised using a sterile razor blade, and placed into 40 mL of seawater from the lab’s inflow system. The tissue was then homogenized in a sterilized mortar and pestle for 10 min. Half of the tissue homogenate was treated with 10,000U of proteinase k (Sigma-Aldrich) and incubated for 1 h at 37°C. Two *Pisaster ochraceus* were inoculated with 10 mL of crude tissue homogenate and two *P. ochraceus* inoculated with the proteinase-k treated homogenate by direct injection into their coelomic cavity. The inoculated asteroids were then placed in individual plastic flow-through aquaria (under low-flow conditions). Individuals were monitored per the flow-rate experiments described above.

### Comparison of wasting time (lesion genesis) with experimental parameters

We modeled the time to wasting (i.e. lesion genesis time) across all experiments against available parameters, which varied between experiments (Table S2). All statistical analyses were performed using XLStat version 2019.4.1 in Microsoft Excel. Response of lesion time to treatment in OM addition experiments were examined by least squares mean ANOVA. The relationship between lesion time and comparison variables was performed by multiple linear regression, forward or backward selection procedure and change in Akaike’s AIC as entry criterion.

### Association of wasting susceptibility with rugosity and surface area to volume ratio

We investigated whether inherent asteroid properties related to wasting susceptibility in context of BLODL by examining their rugosity (i.e. corrugated-ness), which is a key determinant of boundary layer extent. Individual, intact whole-animal specimens of asteroids were collected from several locations (Table S3) and immediately preserved in 20% neutral buffered formalin. All individuals were transported to the lab at Cornell University. Computed tomography was performed on whole specimens at the Cornell University Equine Hospital without contrast to estimate surface area: volume using a Toshiba Aquillon computed tomographic multi-slice scanner.

The relative rugosity between wasting-affected and less wasting-affected asteroid species was examined by calculating the ratio of 3D (determined by computed tomography) to 2D (as calculated below) for each asteroid specimen and comparing between species. Asteroid species were categorized based on prevalence of wasting (less or not affected = *Dermasterias imbricata*, *Henricia leviuscula*, *Patiria miniata*; wasting affected = *Pisaster ochraceus*, *Solaster stimpsoni*, *Pycnopodia helianthoides*, *Leptasterias* sp., *Asterias forbesi*, *Orthasterias kohleri*, *Pisaster giganteus* and *Pisaster brevispinus*) as reported elsewhere (Montecino-Latorre et al., 2016;Bucci et al., 2017;Miner et al., 2018;Jaffe et al., 2019;Konar et al., 2019). These were likewise compared between animal volume, surface area:volume and 2D area.

Because the resolution of computed tomography is only 400 µm, which is larger than some surface features (e.g. papulae), we performed micro-CT (µCT) analyses on one large and one small individual of several key species after staining for at least 24 h in IKI solution (Fig. S4). X-Ray µCT data were analyzed using the Avizo version 2019.4 software (ThermoFisher Scientific). Briefly, 2-D image slices were uploaded and stacked to reconstruct a 3-D volume for each specimen. A median filter was applied to each 3-D reconstruction to reduce noise and smooth edges. The volume of interest was isolated from surrounding background using a thresholding approach. Next, the total volume for each specimen was segmented into 1 cm sections (each composed of 500 stacked 20 µm slices). After eliminating holes from each 1 cm sub-volume, the total volume, surface area, and rugosity were determined using the “Label Analysis” module of the Avizo software. The surface areas reported were calculated by subtracting the 2-D surface areas of the two, flat end slices from the total 3-D surface area for each 1-cm segment. The relationship between ray length and surface area was investigated by linear regression. We first calculated total two-dimensional area for each specimen by taking into consideration central disc radius and assuming triangular shape of rays, accounting for total height of central disc and height of ray tip (see Fig. S5). We then used the ratio of total surface area (determined by CT) to calculate surface area as a measure of rugosity. All image stacks for micro-CT analysis are available from MorphoSource (Duke University) under project accession P1047.

### Asteroid Specimen Respiration

We hypothesized that specimens with greater respiratory O_2_ demand relative to their calculated O_2_ flux would be more susceptible to wasting. The respiration rates of individual asteroids was measured upon experiment initiation for *Asterias forbesi* and *Pisaster ochraceus*, and for additional species at the Bodega Marine Laboratory (Table S4). HOBO O_2_ Loggers (U26; Onset) were placed into sealable plastic containers to which individuals were added. The incubations were circulated using a battery-operated submersible DC motor and propeller. The containers were then filled by immersion in flow-through seawater, and sealed, excluding all visible bubbles. Individuals were incubated for 1 – 2 hr in containers before retrieval of O_2_ probe. Respiration rate was calculated by the linear change in O_2_ concentration over time in incubations. Respiration rates were compared to calculated maximum diffusion rates based on overall surface area determined by computed tomography using Fick’s second law of diffusion (J = -D*∂C/∂d where J = flux across the membrane, D = diffusivity constant of O_2_ in seawater, C = concentration difference between coelomic fluid and seawater – in this case assuming completely anoxic coelomic fluid and saturated seawater, and d = thickness of outer epithelium-assumed to be 20 µm.)

### Time series analyses of wasting intensity and chlorophyll a at Whidbey Island

To understand the relationship between primary producer biomass (chlorophyll a), physico-chemical parameters (temperature, salinity, dissolved O_2_), and occurrence of wasting, we examined data obtained from the Penn Cove Shellfish data buoy and compared this to observations of wasting frequency at Coupeville Wharf and Langley Harbor as reported previously (Hewson et al., 2018) from August 2014 to June 2019 (i.e. 5 years). We also compared wasting frequency with precipitation data obtained from the National Center for Environmental Information (NOAA), which may be seen as a proxy for potential terrestrial runoff. We first calculated the mean time of wasting over the 5-year period and compared this to 5-year mean values of all parameters. We then performed at-time-of-wasting to prior to wasting comparison following a shifting window approach comparing the 3-month window immediately before wasting with 3-month windows in earlier months (Fig. S6).

### Stable isotopic signatures in historical wasting asteroid specimens

The natural abundance of ^15^N and ^13^C was determined in 71 individual starfish specimens, including 50 individuals representing paired asymptomatic/wasting affected species at distinct sites and sampling times which were collected as part of prior work (Hewson et al., 2014) (Table S5). We included an additional 21 individuals of different species to provide context of stable isotopic composition. Samples were collected and frozen at −20°C prior to analysis. Thawed tissue samples were subsectioned for analysis by scraping tube feet into sterile 1.2 ml cryovials. Samples were freeze-dried at −45° C for one week then ground with mortar and pestle. A subsample of 1 mg of tissue was encapsulated into tin and subsequently analyzed on a Carlo Erba NC2500 Elemental Analyzer coupled to a Thermo Scientific Delta V Advantage IRMS (Bremen, Germany).

## RESULTS AND DISCUSSION

The results of our work provide support for our hypothesis that sea star wasting is associated with the formation of anaerobic conditions adjacent to asteroid surfaces. First, we show that organic matter amendment leads to faster lesion genesis than untreated stars, which is preceded by increased bacterial abundance in some treatments and the proliferation of copiotrophic taxa on and above asteroid surfaces in all treatments. In asteroids which wasted in the absence of external stimuli, epidermal and body wall tissues showed a similar progression of copiotrophic bacterial orders, and at the time of lesion and until death the proliferation of strict and facultative anaerobes. Next, we show that lesions form as a consequence of exposure to suboxic water column conditions. We also demonstrate that wasting is correlated with primary production trends over 5 years at a field site. Finally, we provide further evidence of predominantly anaerobic conditions during mass mortality in 2013 – 2014 by way of enriched ^15^N pools in affected tissues relative to asymptomatic individuals.

### Organic matter amendment stimulates boundary layer microorganisms and results in rapid wasting

We sought to examine the impact of elevated heterotrophic bacterial respiration on animal surfaces through amendment with various sources of OM which we hypothesized would fuel microbial remineralization. We performed laboratory experiments in which *P. ochraceus* was amended with peptone, *Dunaliella tertiolecta*-derived particulate OM (POM), and coastal seawater POM and examined their impacts on SSW progression and boundary layer bacterial abundance and composition. The addition of organic substrates (peptone and *Dunaliella tertiolecta*-derived POM) induced significantly faster lesion genesis than control incubations (p=0.012 for peptone and p=0.04 for *Dunaliella*-POM, Student’s t-test, df=5), but lesion genesis time was not significantly different for the addition of coastal-POM (Fig. 2). Collective treatment temporal pattern of lesion genesis was only significantly different from controls with amendment with peptone (p = 0.0154, log-rank test, df=5) and *Dunaliella*-POM (p=0.0339, log-rank test, df=5). Variation in dissolved O_2_ in incubations varied over the course of the experiment from 9.6 – 10.2 mg L^-1^ and were never under-saturated. Temperature varied from 12 – 14°C, but variation did not correspond with wasting in any treatment.

**Fig 2:**
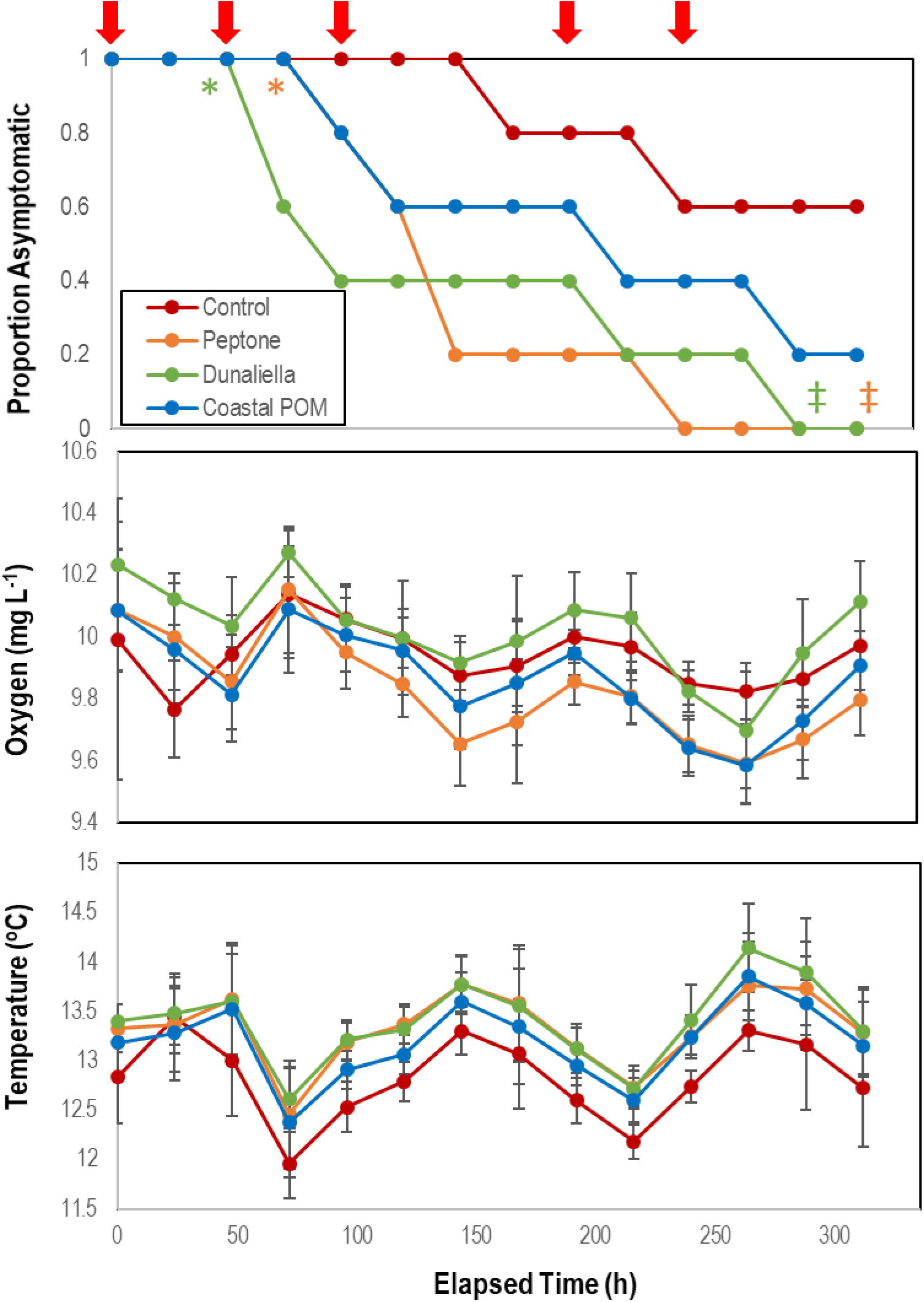
Proportion of asymptomatic *P. ochraceus* (n = 5 each treatment) incubated in flow-through conditions at the Bodega Marine Lab in response to organic matter enrichments (Peptone, *Dunaliella tertiolecta* culture POM, and coastal POM collected from the inflow at the Bodega Marine Laboratory in August 2019. Dissolved O_2_ and temperature were measured in flow-through sea tables bearing each OM treatment. Error Bars = SD. The red arrows above the top panel indicate sampling for microbiome analyses. * indicates wasting speed (i.e. time to appearance of first lesion) was significantly (p<0.05, Student’s t-test) faster than control. ‡ indicates that the the overall trend in lesion formation was significantly different to controls (p< 0.05, log-rank test).

The boundary layer microbiota of *P. ochraceus* during the organic matter amendment experiment changed over time in all treatments (Figs. 3E-G; Figs S7), but the most prominent changes were distinguished by the copiotrophic orders Flavobacteriales and Rhodobacterales (Flavobacteriales; control: p < 0.001, *Dunaliella*: p = 0.002, peptone: p = 0.001, coastal POM: p < 0.001, ANOVA fit with a Generalized Linear Model) (Rhodobacteriales; control: p < 0.001, *Dunaliella*: p < 0.001, peptone: p = 0.010, coastal POM: p < 0.001, ANOVA with a Generalized Linear Model), which increased uniformly in all incubations, indicating that captivity alone may stimulate these groups (i.e. containment affect; Fig. 4). We also observed evidence for treatment-specific proliferation of genera with OM enrichment. Unamended and peptone supplemented *P. ochraceus* experienced the most consistent change in Flavobacteriales, with both conditions exhibiting a linear increase in population mean relative to the mean of all other sub-OTUs. Flavobacteriales in coastal POM-supplemented *P. ochraceus* were elevated from the first to final timepoints, but were primarily distinguished by a large boom and bust after 96 h (c.f. bacterial abundance below). This spike was due to an increase in the family *Crocinitomicaceae*, which, in addition to the family *Flavobacteriaceae*, comprised the majority of Flavobacteriales across all treatments. Rhodobacterales, which primarily consisted of the family *Rhodobacteraceae*, increased in all experimental conditions.

**Fig. 3:**
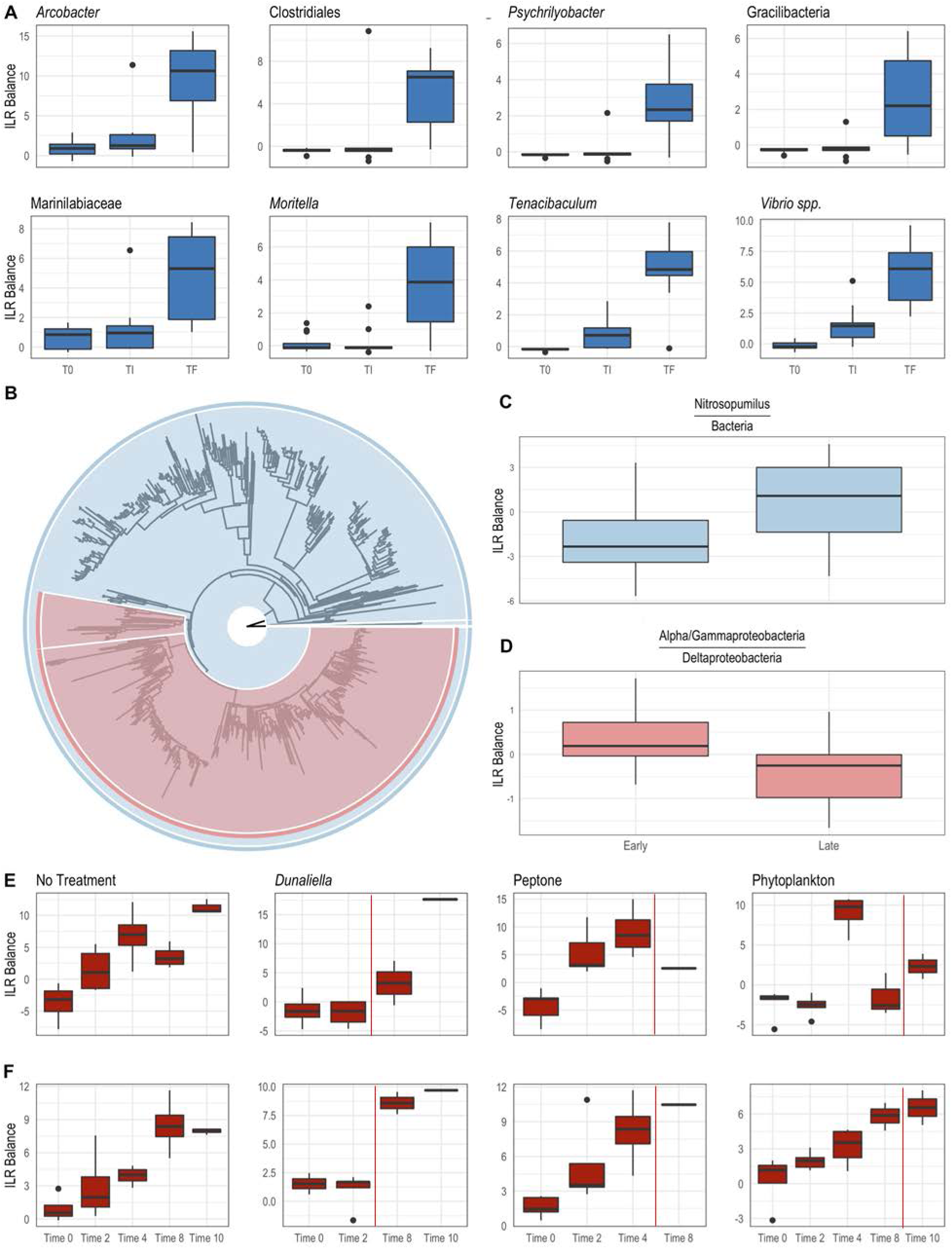
Differential abundance of bacterial taxa from body wall samples (A; *P. ochraceus* June 2018) and surface swabs (B-F; *P. ochraceus* August 2019). (A) Boxplots were derived using PhyloFactor (Washburne et al. 2017), which uses a generalized linear model to regress the isometric log-ratio (ILR balance) between opposing clades (contrasted by an edge) on a phylogenetic tree. This was done iteratively, with each iteration, or factor, maximizing the F statistic from regression. Shown taxa represent either a single factor or combination of factors (when, for example, multiple factors identified different sOTUs with the same taxonomic classification). Labels represent either the highest taxonomic resolution or the highest classification shared by all sOTUs of a given clade. T0 = experiment commencement, TI = lesion genesis, TF = time of death. (B-D) Balance contrast of early (before lesion genesis) samples compared to late (immediately prior to lesion genesis) samples. Samples were transformed using the Phylogenetic Isometric Log-Ratio (PhILR; Silverman et al. 2017) transform, which uses a phylogenetic tree (B) to convert an sOTU table into a new matrix of coordinates derived from the ILR of clades that descend from a common node. We used a sparse logistic regression with an *l*_1_ penalty of λ=0.15 (Silverman et al. 2017) to analyze the ILR at each node, and included a select number of ‘balances’ with positive coefficients (C-D). (C) is the balance of *Nitrosopumilus* (colored blue in (B), comprises the thin sliver on the right side of the tree) relative to the rest of the dataset (also shown in blue in (B)). A positive shift indicates an increase in *Nitrosopumilus* relative to its denominator. (D) is the balance between a clade of Alpha/Gammaproteobacteria (large, red clade in (B)) and Deltaproteobacteria (Bdellovibrionales and Desulfobacterales; small, red clade in (B)). A negative shift indicates that the denominator, Deltaproteobacteria, is increasing relative to Alpha/Gammaproteobacteria. (E) and (F) were derived from a PhyloFactor object and show the ILR balance of Flavobacteriales (E) and Rhodobacterales (F) relative to all other sOTUs. Organic amendment is given above boxplots. Time 0 = 0 h, Time 2 = 48 h, Time 4 = 96 h; Time 8 = 192 h; Time 10 = 240 h. Total sample numbers for each treatment (which varied due to the loss of asteroids over the course of the experiment to wasting) is given in Fig. 4. The vertical red line in panels E and F indicate the average time at which asteroids formed lesions.

**Fig 4:**
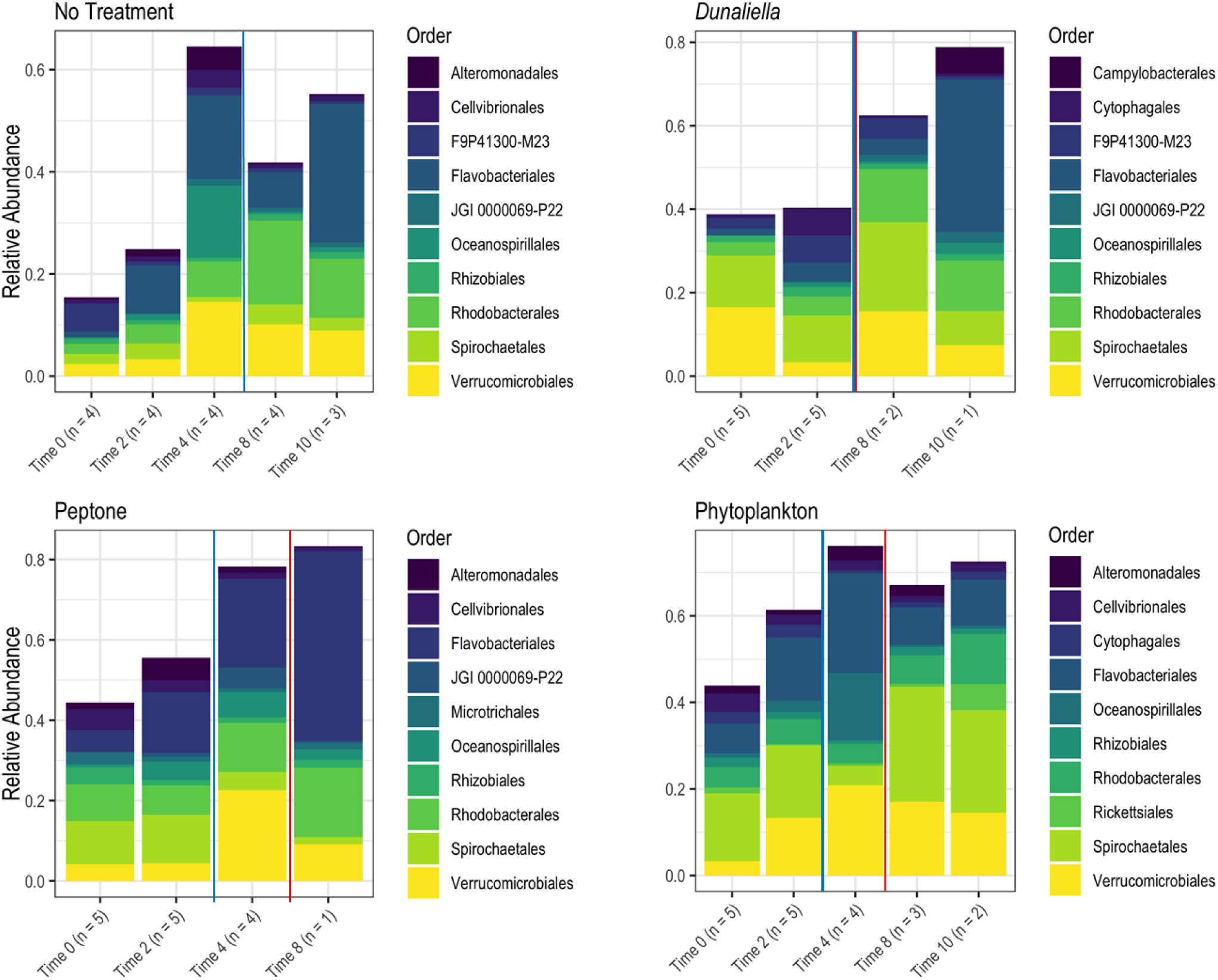
Relative abundance of bacterial orders derived from *P. ochraceus* epidermal swabs. Specimens were enriched with the indicated organic material and sampled until lesion genesis. Time 0 represents initial sampling and each subsequent time indicates the respective day. n values reflect the number of healthy specimens at each given timepoint. The solid blue line on each panel indicates when lesions first formed per treatment, and the solid red line on each panel indicates the mean lesion time within the treatment. Time 0 = 0 h, Time 2 = 48 h, Time 4 = 96 h; Time 8 = 192 h; Time 10 = 240 h.

Bacterial cell abundance on surfaces (relative to both initial values and controls) illustrated large swings prior to wasting onset (Fig. 5). Individuals that did not waste over the course of the experiment maintained abundances of 0.7 – 2.6 x 10^6^ cells mL^-1^, which was enriched 53 to 1743% above bacterioplankton abundances in incubation treatments (Fig. 5). On aggregate, wasting stars had higher bacterial abundances than non-wasting stars (Fig. S8), however the relationship was not significant because of high variation between treatments with OM. Treatment bacterial abundances remained no different to controls over the first 48 h of incubation, but increased relative to controls in peptone and coastal-POM treated asteroids after 72 and 96 h, respectively. However, by 96 h for peptone and 120 h for coastal POM both amendments had again declined, and remained no different to controls after this time (Fig. 5). In contrast, bacterial abundance in *Dunaliella*-POM incubations were no different to controls over the first 48 h of incubation, and were far less than controls after 72 h. Since bacterial abundance increased prior to lesion genesis in at least two OM treatments, we posit that wasting is influenced by copiotroph proliferation on animal surfaces. The decrease in bacterial abundance after initial increase in both peptone, coastal POM, and consistently lower bacterial abundance in *Dunaliella*-POM incubations may be evidence of heterotrophic remineralization-fueled O_2_ deficit over time on wasting asteroids, similar to the effect observed in our experiments with *Asterias forbesi* incubated in hypoxic water (see below). Facultative and strict anaerobes generally experience slow growth rates compared to aerobic taxa because it is less energetically efficient to grow on reduced electron acceptors. While standing stock of aquatic bacteria may be higher in anaerobic conditions than in aerobic conditions, population growth rates are typically lower (Cole and Pace, 1995).

**Fig 5:**
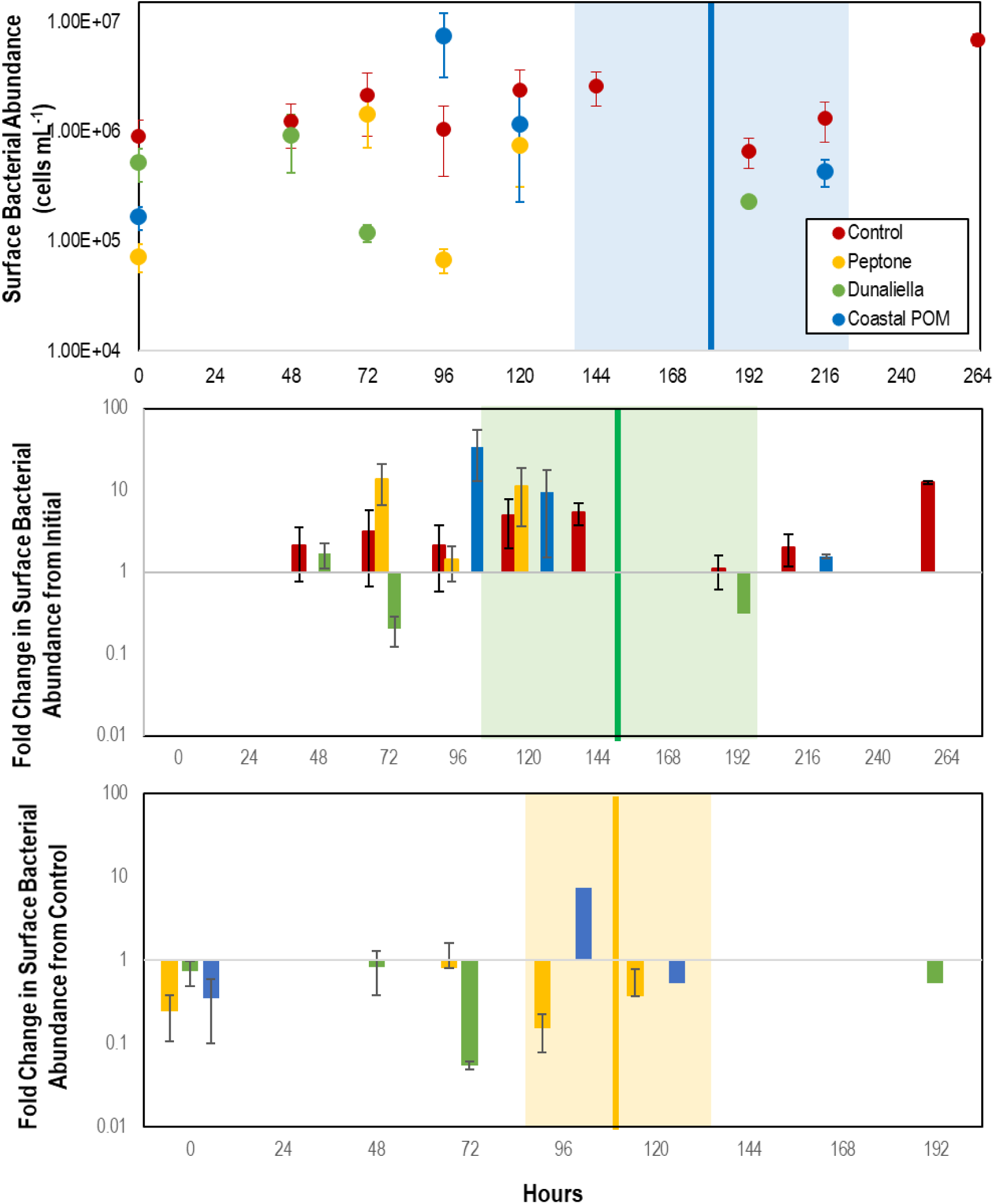
Abundance of bacteria in proximity to *P. ochraceus* surfaces (top), percentage change from initial (middle), and relative to controls (bottom). during first 10 days of experiment in response to organic matter enrichment (n = 5 for each treatment) as assessed by SYBR Gold staining and epifluorescence microscopy. Control specimens are indicated in red, while the mean of stars that wasted in Peptone, *Dunaliella tertiolecta* POM, and Coastal POM are indicated separately. The solid vertical line on the top panel represents the mean time that asteroids developed lesions in the coastal POM treatment, the solid line on the middle panel represents the mean time for lesion development in *Dunaliella tertiolecta* POM treatments, and solid line on the bottom panel represents the mean time for lesion development in peptone treatments (separated between panels for clarity). The shaded regions represent lesion development standard error for respective treatments.

### Shifts in heterotrophic bacterial and archaeal communities during wasting progression in the absence of external stimuli

Because sampling by biopsy punch imparts stress on animals that may elicit wasting, sampling in the absence of external stimuli focused on samples collected at the time of lesion genesis (August 2018) and samples collected after lesion genesis (May – June 2018), and are distinct from samples collected by surface swab (collected in organic matter enrichment experiment). Hence, compositional changes observed in these surveys likely reflect a combination of taxa that change prior to lesion formation and those that degrade tissues.

In the August 2018 study, *Pisaster ochraceus* developed lesions without stimuli beginning 5 d after isolation, and by 8 d more than half of incubated stars were symptomatic (Fig. 6). Lesions formed initially concomitant with an approx. 2°C swing in temperature, however continued in other stars as temperatures progressively decreased over the course of the experiment. Initial genesis of lesions was not accompanied by variation in either DO or pH. Lesions were grossly characterized by nonfocal loss of epidermal tissues, which exposed underlying body wall tissues. Lesion margins were not remarkable in terms of coloration which would otherwise indicate melanization. Epidermal samples from these specimens revealed no significant difference in microbial composition between artificial and natural lesions in 3 specimens that wasted (PERMANOVA; P = 0.139; Fig. S9). Between initial samples and the time of lesion genesis, most taxa identified as differentially abundant were less abundant sub-OTUs. In the May-June 2018 study, we observed a progressive increase in copiotrophic orders between initial samples and those taken shortly after lesions had formed, including Campylobacterales (p<0.001), Flavobacteriales (p<0.001) and Vibrionales (p<0.001; ANOVA fit with a Generalized Linear Model) (Fig. 3A). This occurred concomitant with an increase in *Nitrosopumilus* and obligate anaerobes (Deltaproteobacteria) relative to a large clade of typically fast-growing phyla (Alpha- and Gammaproteobacteria) (Fig. 3B-D).

**Fig. 6:**
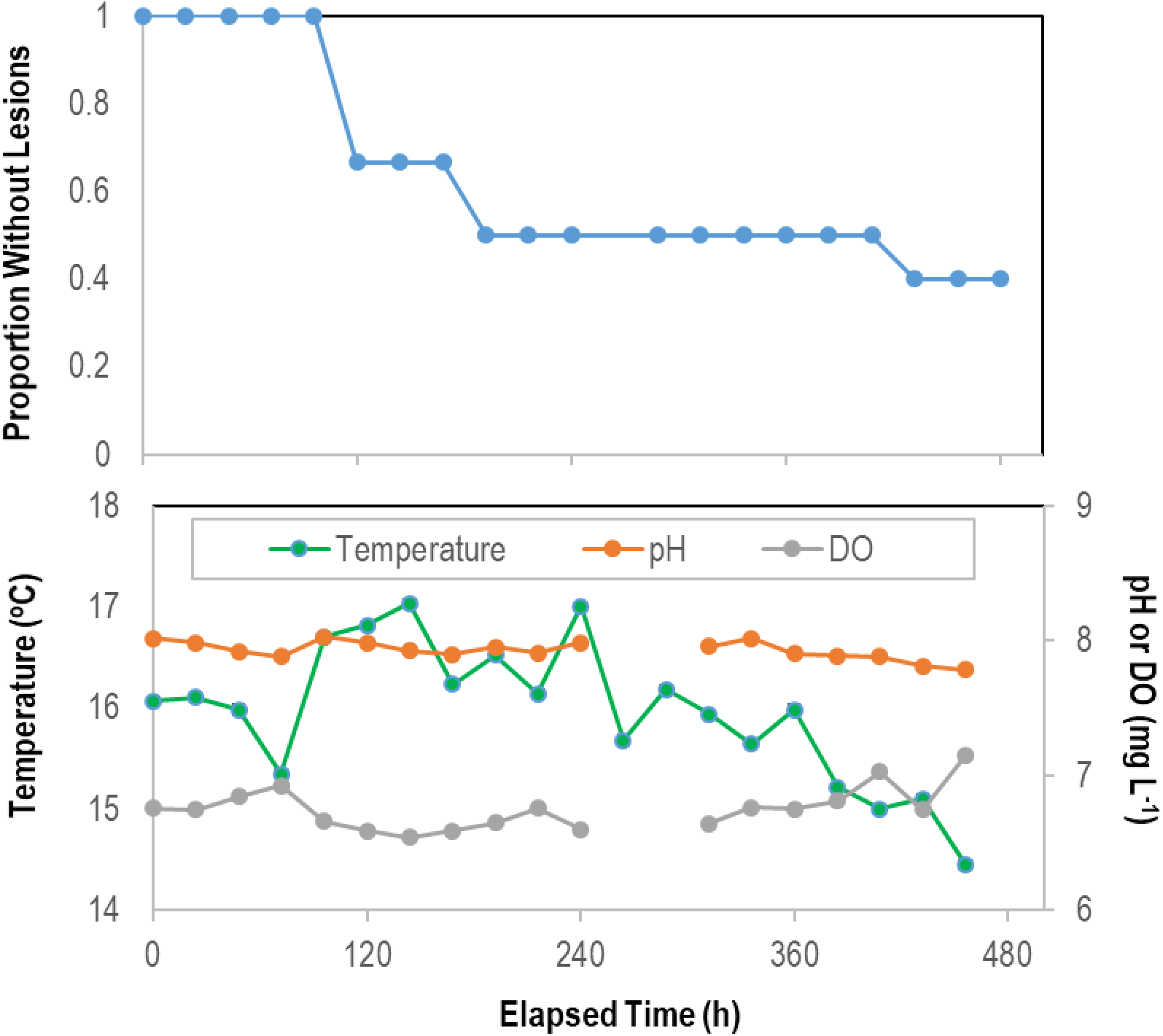
Proportion of asymptomatic *P. ochraceus* (n = 5) remaining over time during longitudinal study of microbiome composition in the absence of external stimuli. The mean flow rate into aquariums was 3.81 ± 0.05 mL s^-1^ (average residence time in aquariums 37 min).

Following lesion formation, bacteria on surfaces likely experience a complex milieu of OM molecules, including those from decaying tissues. We found evidence for the proliferation of anaerobic taxa under these conditions by comparing microbial composition between lesion formation and animal death. Between lesion genesis and animal death, we observed a further increase in copiotrophs in body wall (May-June 2018) and epidermal (August 2018) samples, and a proliferation in microaerophiles (*Arcobacter* spp.), facultative anaerobes (*Moritella* spp.), and obligate anaerobic families (Clostridia, Fusobacteria and Bacteroidia) at time of death (Fig. 3A; Fig. S10). The two sub-OTUs with the largest F-statistic from regression over the entire course of wasting from initial samples to animal death belonged to the families *Desulfobulbaceae* (p<0.001) and *Desulfovibrionacea* (p=0.002; ANOVA fit with a Generalized Linear Model). Both of these families are strictly anaerobic sulfate reducers. These results are consistent with the pattern of microbial assemblage variation observed in the organic matter amendment experiment. Previous study comparing wasting (specimens already had lesions) and asymptomatic asteroid-associated community gene transcription also noted the increase in transcripts from *Propionibacterium*, *Lachnospiraceae* and *Methanosarcina*, which are strict anaerobes, as well as *Stigmatella* and *Staphylococcus*, which are facultative anaerobes, as a proportion of total transcripts (Gudenkauf and Hewson, 2015).

### Microbial assemblage composition and abundance variation illustrates an increasingly anaerobic environment near asteroid surfaces

Taken together, these studies suggest that lesion formation is associated with a general proliferation of bacteria, including well-known copiotrophic orders (including *Flavobacteriaceae* and *Vibrio* spp.) of marine bacteria. Several studies have observed the proliferation of copiotrophic taxa longitudinally during wasting, including genera within the families Flavobacteriaceae, Rhodobacteriacaea (Lloyd and Pespeni, 2018), Actinobacteria, and genera in the orders Altermonadales (Nunez-Pons et al., 2018), Vibrionales and Oceanospiralles (Hoj et al., 2018). These taxonomic groups are amongst the most active constituents of bacterioplankton and major players in marine OM degradation, some of which have facultative anaerobic metabolisms (Pinhassi et al., 2004;Choi et al., 2010;Buchan et al., 2014;Thiele et al., 2017;Pohlner et al., 2019). While it is tempting to ascribe pathogenicity traits to groups that are enriched on disease-affected tissues (based on members of the same family or genus causing pathology), or infer their role in community dysbiosis (i.e. the microbial boundary effect), this is not possible in the absence of demonstrated pathogenicity or strain-level assignment (Hewson, 2019). The observation of strict anaerobic bacteria in underlying tissues after lesions had formed suggest the creation of a depleted oxygen environment on asteroid surfaces in response to organic matter loading.

All aquatic surfaces are coated with a thin film of water (i.e. diffusive boundary layer) that impedes gas and solute exchange, and, provided aerobic respiration is sufficiently high, suboxic conditions can form on a surface despite oxygen saturated water circulating above (Jørgensen and Revsbech, 1985). This may result in the proliferation of facultative and obligate anaerobes until asteroid death. Stimulation of bacteria and subsequent O_2_ diffusion limitation is well described in mammalian respiratory systems as well, and is especially pronounced in cystic fibrosis patients. Heterotrophic bacteria inhabiting mammalian lungs thrive on mucins and generate biofilms which further restrict O_2_ diffusion into tissues. O_2_ consumption by biofilms and by neutrophils may result in hypoxia and reduced diffusion of O_2_ across alveolar tissues (Wu et al., 2018). This in turn leads to the proliferation of anaerobes, which are present in clinically normal lungs (reviewed in Guilloux et al., 2018) and elevated in diseased lungs (Denner et al., 2016;Wang et al., 2019;Spence et al., 2020). This phenomenon is also observed in fish gills (Legrand et al., 2018;Meyer et al., 2019) which are inhabited by copiotrophic and potentially facultatively anaerobic taxa (Reverter et al., 2017;Rosado et al., 2019).

Rapid mineralization of dissolved OM by bacteria near asteroids has been noted previously in studies of epidermal amino acid uptake by *Asterias rubrens*, which ultimately led to decreased animal weight (Siebers, 1979), and may be responsible for very low ambient DOM concentrations adjacent to asteroids (Siebers, 2015). Remineralization of OM by heterotrophic microorganisms fuels oxygen consumption, which may in turn lead to oxygen deficit when consumption is not matched by gas diffusion. For example, excess phytoplankton-fueled bacterial respiration, caused by eutrophication and enrichment from terrestrial sources and upwelling zones may result in ‘dead zones’ (e.g. Mississippi River Plume, Peruvian upwelling zone, Benguela current; reviewed in (Diaz and Rosenberg, 2008) and may be exacerbated by seasonal temperature changes (Murphy et al., 2011) and restricted bathymetry (Diaz, 2001). We posit that OM amendment stimulates bacterial abundance immediately adjacent to asteroid respiratory surfaces (i.e. within boundary layers) leading to suboxic microzones and ultimately limiting gas diffusion potential (Gregg et al., 2013).

Bacterial stimulation and enhanced wasting in asteroids is paralleled by the DDAM (dissolved organic carbon, disease, algae, microorganism) positive feedback loop in tropical corals (Dinsdale et al., 2008;Barott and Rohwer, 2012;Silveira et al., 2019). Coral disease is associated with OM enrichment (David et al., 2006;Smith et al., 2006), some of which originates from sympatric primary producers (Haas et al., 2010;Haas et al., 2011), which in turn are more labile than OM released from the corals themselves (Haas et al., 2016;Nakajima et al., 2018) and results in both elevated bacterial abundance on coral surfaces (Dinsdale and Rohwer, 2011;Haas et al., 2016), and enhanced remineralization rates (Haas et al., 2016). Bacteria at the coral-water interface have higher energetic demands than those in plankton (Roach et al., 2017), and are highly adapted to organic carbon availability in their local environment (Kelly et al., 2014). The spatial scale on which bacteria react to OM is primarily at water-surface interfaces (Brocke et al., 2015). Hypotheses for the mechanism of coral mortality caused by heterotrophic bacteria include disruption in the balance between corals and their associated microbiota (David et al., 2006), introduction of pathogens that have reservoirs on macroalgae (Nugues et al., 2004), or dysbiosis resulting in invasion by opportunistic pathogens (Barott and Rohwer, 2012). In black band disease, DOC released from primary production causes micro-zones of hypoxia which result in production of toxic sulfides, which in turn result in opening of niches for cyanobacteria (Sato et al., 2017). In asteroid wasting, the proliferation of heterotrophic bacteria and wasting disease may be due to any of these effects.

### Asteroid wasting is induced by suboxic conditions

Our data demonstrate that SSW is induced by suboxic water column conditions. We incubated *A. forbesi* in suboxic water and observed patterns of wasting progression, boundary layer bacterial abundance and microbial assemblage β-diversity. Dissolved oxygen (DO) concentrations were controlled in an aquarium setting by continuous sparging with N_2_, which were on average 39% lower than untreated control incubations (Fig. 7). All individuals remained asymptomatic in control incubations over the 13 day experiment, while 75% of individuals in hypoxic conditions developed lesions (mean time to lesion genesis = 9.58 ± 0.89 d; Fig. 7). Development of lesions over time was strongly related to treatment (p = 0.006, log-rank test, df=12). Bacterial abundance on animal surfaces (which we define as abundance in surface samples) corrected for aquarium water values increased in both control and suboxic treatments over the first 6d of incubation, but by day 13, abundance of bacteria in suboxic treatments was significantly lower (p<0.001, Student’s t-test, df=12) on suboxic treated individuals than in control individuals (Fig. S11). *Asterias forbesi* treated with suboxic waters demonstrated consistent shifts in microbial communities with treatment (Fig. S9). However, no single bacterial taxonomic organization strongly differentiated normoxic from suboxic conditions.

**Fig. 7:**
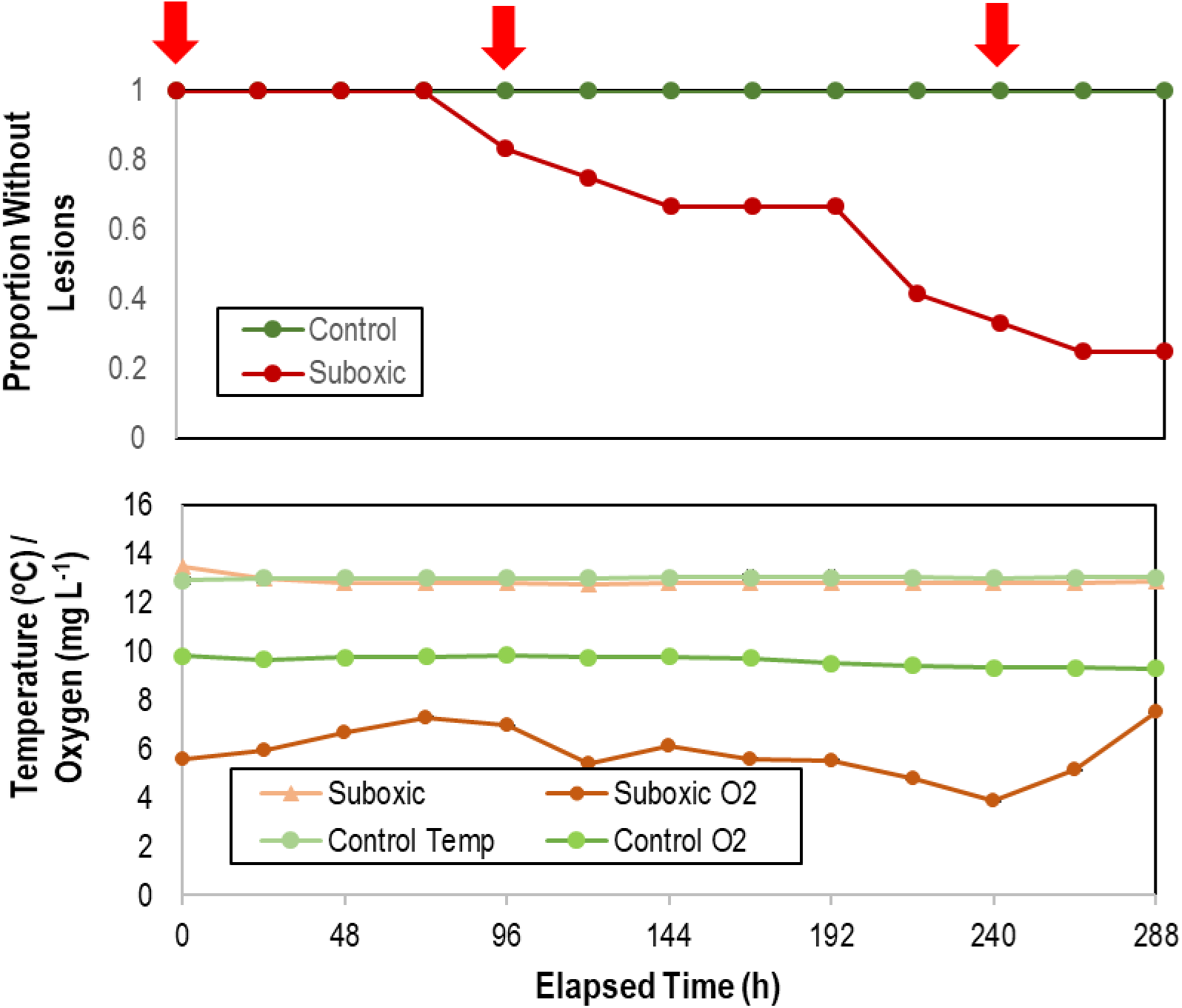
Proportion of asymptomatic *Asterias forbesi* incubated in normoxic and suboxic water (top) and variation in temperature and O_2_ in incubation aquaria (bottom) over time. The red arrows at top indicate samples which were included in analysis of microbiome composition (see Fig. S9).

Further evidence for the role of oxygen in SSW was observed in experiments comparing wasting speed under variable incubation flow rates (i.e. water replenishment rates). While we did nt measure DO concentrations in these experiments, higher flow rates likely had higher DO concentrations than lower flow rates, in addition to reducing OM (e.g. mucus) and toxic exudates in animal waste (notably NH_3_ [Propp et al., 1983] and S^-^ [Vistisen and Vismann, 1997]). Furthermore, higher flow rates may have experienced less extensive boundary layers than lower flow rates (Fonseca and Kenworthy, 1987). The time to lesion genesis in *Pisaster ochraceus* was faster for asteroids under low-flow conditions than those under high flow conditions (p = 0.006, Student’s t-test, df=3), however survival was not significantly different between flows (log-rank test, ns; Table S6). Desiccation, which was used to insult asteroids and simulate emersion during low tide events in warmer temperatures, resulted in faster lesion formation (p = 0.05, Student’s t-test, df=3) under low flow conditions, but not different under high flow conditions (log-rank test, ns; Fig. 8). Addition of tissue homogenates, which was initially used to assess transmissibility of tissue-derived agents, resulted in faster wasting than addition of proteinase k treated tissue homogenates (p = 0.04, Student’s t-test, df=3); however, survival was no different between control treatment low flow and the addition of proteinase K-treated or untreated tissue homogenates (log rank test, ns; Fig. 8).

**Fig. 8:**
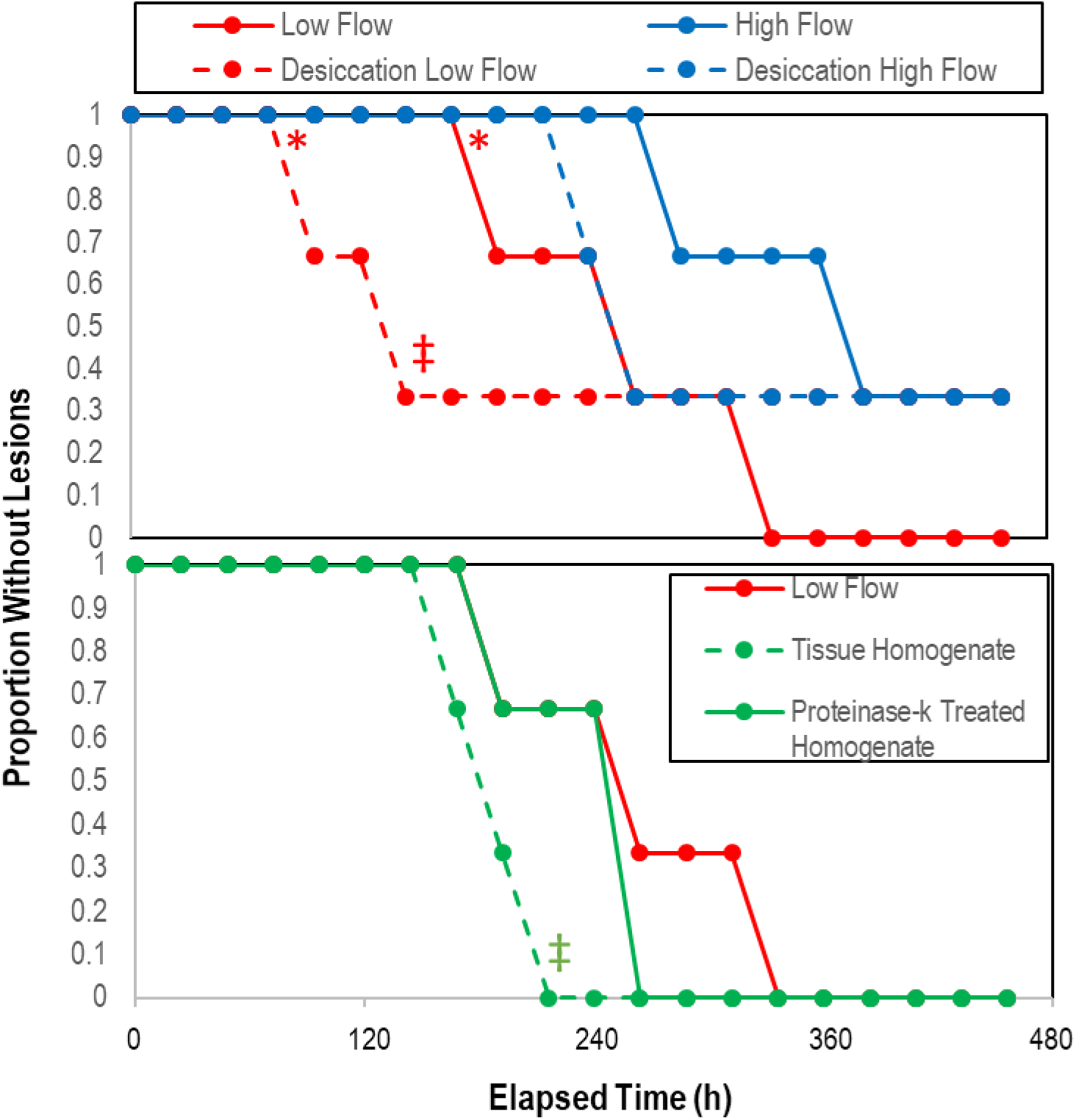
Proportion asymptomatic *P. ochraceus* over time in response to treatment (top) desiccation (n = 3), and (bottom) treatment with crude and proteinase k-treated tissue homogenates (n = 2). * indicates that low flow lesion genesis time was significantly (p<0.05, Student’s t-test) compared to high flow rate; ‡ indicates that the overall trend of desiccation under low vs high flow rate and with the addition of proteinase-k treated homogenate vs low flow controls was significant (p< 0.05, log-rank test).

The mechanism by which asteroids are particularly sensitive to ambient O_2_ concentrations is not well constrained by empirical studies, especially as it relates to SSW. Asteroids mostly rely on passive respiration (c.f. ventilation) and gas diffusion across outer membranes to meet respiratory demand, a point illustrated by mass mortality events of benthic invertebrates, including asteroids, correlated to low O_2_ conditions (reviewed in Diaz and Rosenberg, 1995;Levin, 2003;Levin et al., 2009). Together, these data point to significant influence of O_2_ conditions on asteroid wasting. While water column hypoxia events were not observed in concert with SSW in 2013 and beyond, spatially localized hypoxia may occur near surfaces experiencing limited hydrodynamic flow (Gregg et al., 2013).

Analysis of covariance revealed that lesion time was explained best by different parameters, depending on the experiment. For *Asterias forbesi*, lesion genesis time variation was best explained by both overall animal mass change during the experiment and initial bacterial abundance (p = 0.12), while in *Pisaster ochraceus* experiments, both OM addition and combined with flow rate was best explained by initial animal mass (p=0.006). In OM addition, variation in lesion genesis time was also explained by change in bacterial abundance over the first 3 d of incubation (p= 0.018) (Table S7). While these observations are entirely correlative – in other words, cannot reveal causality – they further indicate a key role of heterotrophic bacteria inhabiting the asteroid-seawater interface in wasting.

### Wasting is related to inherent asteroid properties that dictate boundary layer extent, gas diffusion, and respiration

Inter- and intra-species susceptibility to asteroid wasting is extensively recorded in previous study, including a significant and positive relationship between individual size and wasting (Hewson et al., 2014), and shifts in size structure after wasting from larger to smaller individuals of *Pisaster ochraceus*, which was believed to be due to recruitment of juveniles (Bates et al., 2009;Eisenlord et al., 2016;Menge et al., 2016;Kay et al., 2019). Wasting in 2013-2014 affected > 20 species of asteroid (Hewson et al., 2014), however the magnitude of SSW impact varied between species. Comparison of community structure before and after wasting suggests inter-species variability in wasting mortality. Asteriid taxa (*Pycnopodia helianthoides, Pisaster* spp., and *Evasterias troschelii*) experienced considerable declines in the Salish Sea (Montecino-Latorre et al., 2016;Schultz et al., 2016) and Southeast Alaska (Konar et al., 2019), while *Dermasterias imbricata* maintained or increased in abundance after mass mortality (Eckert et al., 1999;Montecino-Latorre et al., 2016;Schultz et al., 2016;Konar et al., 2019). In the Channel Islands, SSW disproportionately affected Asteriid taxa relative to *D. imbricata* and *Patiria miniata* (Eckert et al., 1999). Inter-species differences in wasting intensity have been noted in citizen science data accumulated by MARINe (Miner et al., 2018). The potential causes of inter- and within-species wasting susceptibility remain poorly constrained.

We hypothesized that wasting susceptibility may relate to both inter-species variation in rugosity (i.e. degree of corrugation), which dictates diffusive boundary layer thickness, and intra-species surface area-to-volume ratio, which determines total gas flux potential, which are ultimately reflected in patterns of population change since 2013 (Eckert et al., 1999;Montecino-Latorre et al., 2016). Mean and turbulent flow structure around aquatic animals and plants relates to the mean height, density and shape of structures as they compare to flat surfaces (Koch, 1994;Nepf, 2011;Brodersen et al., 2015). Asteroid surfaces bear numerous spines and processes, including papulae, spines, paxillae and pedicellaria. These structures impart rugosity and thus generate diffusive boundary layers proportional to their relative height under both mean and turbulent flow. For example, the boundary layer height above the urchin *Evechinus chloroticus.* can be 4-5 mm under low (1.5 cm s^-1^) flow conditions, which was approximately 2 – 6 X greater than sympatric macroalgae (Hurd et al., 2011). We speculate that more extensive boundary layers may result in a greater deficit in O_2_ due to entrapment of OM adjacent to animal tissues, and also increase the potential for hypoxia at the animal surface since the effective distance over which O_2_ must diffuse is higher in specimens with greater boundary layer extent. Direct measurement of oxygen concentration in boundary layers as they relate to bacterial remineralization are precluded by the sensitivity of instruments (e.g. microelectrodes) to physical damage in non-immobilized specimens.

To explore the relationship between species rugosity and wasting susceptibility, we examined specimens of similar size (n = 26 individual specimens) representing wasting-affected (n = 3) and less/not affected species (n = 5) using whole-animal computed tomography. CT-derived volume was significantly and positively correlated to overall mass across all specimens (p = 0.00001, R^2^=0.9999). To estimate overall surface area of specimens for which respirometry was measured (which were not imaged by CT), we modeled the surface area to ray length, and found it followed an exponential function (mean ray length = 0.0825e^1.0536*LOG(Surface area)^, R^2^ = 0.93. Log (Surface area) was significantly and linearly correlated to Log (Volume; LOG(Volume) = 0.7319*LOG(Surface Area) + 0.7662; R² = 0.9703); Fig. S20). The surface area: volume was significantly and negatively correlated with a logarithmic function defined as SA:Vol = 4.6242e ^-0.549[LOG(Volume)]^) (Fig. S12). The mean rugosity (defined as 3D:2D surface area) was significantly (p = 0.015, Student’s t-test, df = 14) lower in less affected species than more affected species (Fig. 9). Surface area:volume, individual specimen mass, and overall surface area were not significantly different between categories among similarly-sized animals. Because analysis of large animal specimens is limited to a resolution of 400 µm (which is potentially larger than fine-scale features, e.g. papulae on echinoderm surfaces), we performed further analysis on rays of a subset (n = 16) of individuals using micro-computed tomography, which has a resolution of 20 µm. The rugosity of wasting-affected taxa was significantly (p = 0.0002, Student’s t-test, df = 4) greater than less-wasting affected species (Fig. 9). Our observation that more rugose species were more affected by wasting supports the idea that these individuals may be more susceptible because of their greater extent (physical distance) of diffusive boundary layers on respiratory surfaces.

**Fig. 9:**
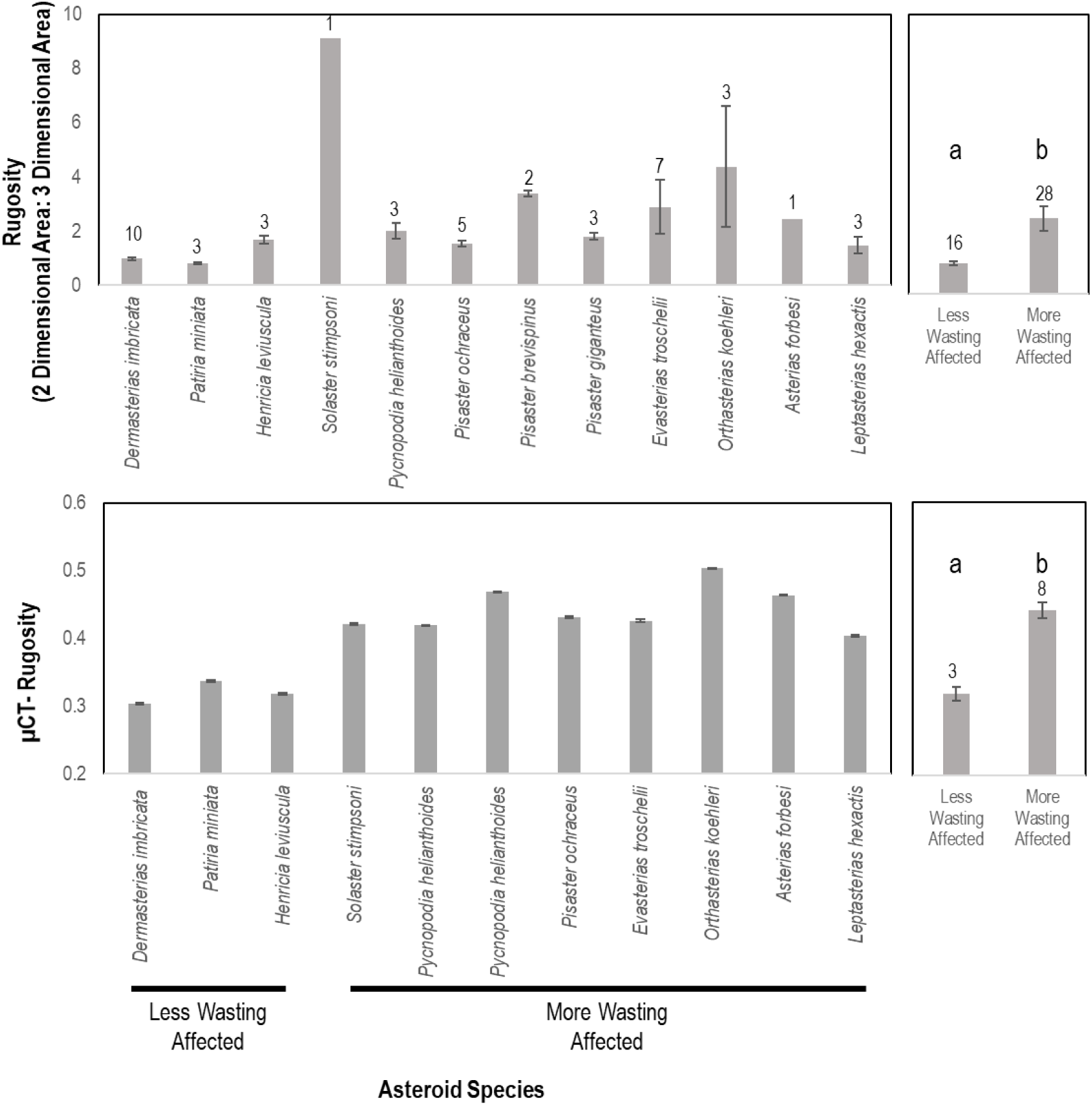
Rugosity of similarly-sized animals between wasting-affected and less wasting affected species as determined by whole animal computed tomography (top) and of an asteroid ray by micro-computed tomography (bottom). a, b denote significant difference at *p* < 0.001. More and less wasting affected were based upon previous work and defined in the text.

Much of the intra-species wasting susceptibility may also be explained by inherent variation in diffusive flux potential. We observed a significant and positive relationship between wasting lesion genesis rate and initial animal mass (p = 0.006; analysis of covariance). Larger individuals have a much lower surface area:volume ratio, where surface area is related to gas flux potential. Under near-surface hypoxic conditions, or when diffusion is impeded by extensive boundary layers, larger individuals are more strongly affected than smaller individuals. We also posit that these observations are the result of more extensive boundary layer height above larger specimens. It is also important to note that ossicle density varies between species (Blowes et al., 2017), and those taxa with lower densities (e.g. *Pycnopodia helianthoides*) were more affected than those with higher densities (e.g. *Pisaster ochraceus*). Species with lower ossicle densities may be differentially susceptible to wasting since their structure may be broken down faster by microbial decomposition or apoptotic processes.

Wasting risk susceptibility may furthermore result from differential diffusive flux potential compared to respiratory demand. We measured the respiration rate (i.e. oxygen demand) of individuals at the start of each experiment, as well as in individuals of several species that were both affected by wasting and those that were less or not affected by wasting that were not a part of experiments to explore whether susceptibility was related to oxygen demand. Mass-normalized measured respiration rates of asteroids were greatest for *Asterias forbesi*, and least for *Dermasterias imbricata* and *Patiria miniata* (Fig. S4). Both *Pisaster ochraceus* and *Asterias forbesi* respiration rates were considerably more than for other specimens. Measured respiration rates for entire animals was compared to theoretical maximum diffusion rates into coelomic fluids (hereafter abbreviated RR:TD). RR:TD was greatest in *Asterias forbesi* and *Pisaster ochraceus* (which were both >1 in most specimens) and least in *Patiria miniata* and *Dermasterias imbricata* (which were always < 0.1). The observed RR:TD corresponds with wasting susceptibility. Perturbation of O_2_ availability in animal surface boundary layers may skew diffusive flux by elongating diffusive path length or reducing differences in O_2_ between tissues and surrounding seawater. Hence, specimens with a higher RR:TD may be more affected by the condition than those with lower RR:TD. We cannot account for variable permeability of outer epidermis between individuals (not measured), and assume that all surface area of asteroids is involved in respiration (which may be over-estimated, since presumably some component of this area comprises mineral structures). Some asteroid species inhabiting typically suboxic environments employ morphological and behavioral strategies to meet O_2_ demand, including nidamental cavities (Johansen and Petersen, 1971;Nance, 1981), cribiform organs (Shick et al., 1981), epiproctoral cones (Shick, 1976), active ventilation of burrows and decreased size of internal organs (Mironov et al., 2016). However, it is unlikely asteroids typically occurring in normoxic intertidal or subtidal conditions have the ability to morphologically adapt to hypoxic conditions.

### Potential sources of OM fueling BLODL

Heterotrophic bacteria in marine environments remineralize OM that originates from autochthonous and allochthonous sources (Ducklow, 1983;Benner et al., 1992;Amon and Benner, 1996). We hypothesize there are two primary sources fueling BLODL: OM from primary production (phytoplankton and macroalgae), and OM from decaying asteroids. Most asteroid wasting is reported in late fall or summer, with fewer reports during other times of the year (Eckert et al., 1999;Bates et al., 2009;Eisenlord et al., 2016;Menge et al., 2016;Montecino-Latorre et al., 2016;Hewson et al., 2018;Miner et al., 2018;Harvell et al., 2019;Hewson et al., 2019). Among the myriad of OM sources in seawater, phytoplankton-derived OM are highly labile (Ochiai et al., 1980;Ogawa and Tanoue, 2003;Thornton, 2014). We propose that wasting is associated with peak or post-peak declines in phytoplankton production in overlying waters, which subsequently results in peak dissolved OM availability. The mean time of wasting mass mortality observed at Langley and Coupeville, Whidbey Island between 2014 and 2019 fell at or within 1 month after the mean annual maximum of chlorophyll a, minimum DO concentration, maximum temperature, and minimum rainfall (Fig. 10). Multiple linear regression (stepwise, backwards selection criteria) revealed a significant model (R^2^=0.866; p=0.001) where temperature (p = 0.006), chlorophyll a (p = 0.027) and salinity (p = 0.044) explained most variation in wasting mass mortality, while forward selection (R^2^=0.774; p=0.0002) revealed that monthly variation in wasting was significantly explained by DO alone. Mass mortality was significantly related (one-way ANOVA, p<0.0001) to elevated chlorophyll in the previous 3 months relative to non-mass mortality months, to elevated salinity, and reduced rainfall (Fig. S6). Because wasting occurs seasonally in late summer and early autumn, correlation between these parameters alone does not necessarily indicate a direct link between primary production and wasting. However, since we also observed that asteroids challenged with a phytoplankton (*Dunalella tertiolecta*) formed lesions faster than unamended controls, it is entirely possible that wasting is related to phytoplankton-derived DOM.

**Fig 10:**
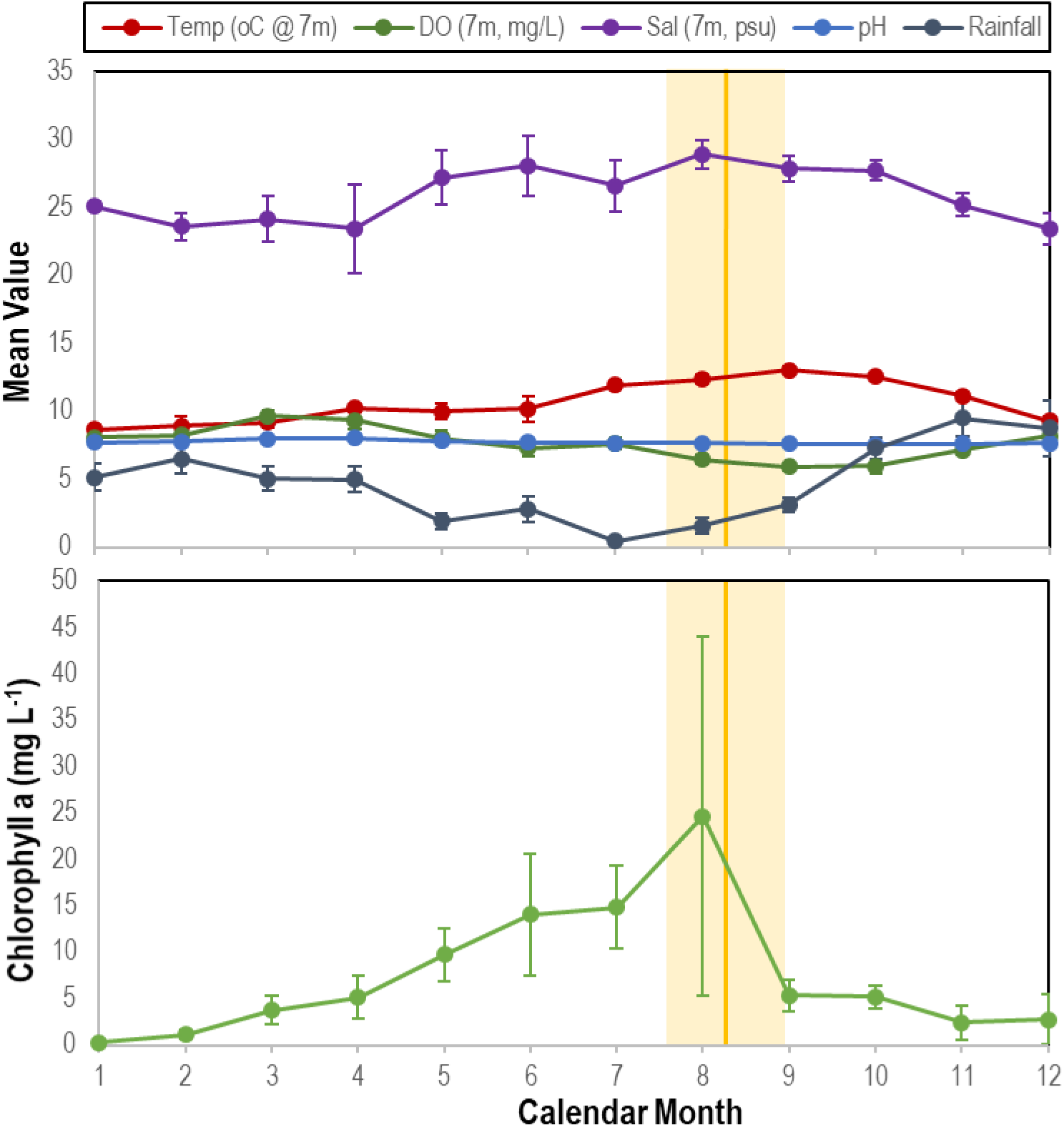
Correspondence between mean time of wasting mass mortality (indicated by solid orange line (SE range indicated by lighter orange bar) compared with physico-chemical parameters (top) and chlorophyll a concentration (bottom) at Penn Cove, Whidbey Island from 2014 - 2019. Temp = temperature; DO = dissolved oxygen; Sal = salinity.

The coherence of wasting with primary production in the Salish Sea raises the question of why wasting mass mortality in the northeast Pacific occurred in the 12 month period following June 2013, especially when asteroids normally persist at sites experiencing very high phytoplankton biomass and only experienced wasting in 2014 (e.g. Cape Perpetua, OR; Leslie et al., 2005). Suchy et al. (2019) observed a prolonged (10 month) period of decreased water column stratification, concomitant with strong predominately southerly winds in fall 2013 and spring 2014 in the northern Strait of Georgia. Chlorophyll a concentrations in the region was also higher in the region in late 2013 compared to the previous 8 years (Suchy et al., 2019) and the 1981-2010 average (Moore et al., 2014) and was marked by a significant peak in late August (later than previous years (Moore et al., 2014), concomitant with the lowest NH_3_ concentration measured over several years at the Seattle Aquarium, presumably a consequence of phytoplankton uptake (Olsen et al. 2016). Mean monthly precipitation was anomalously lower in mid-summer compared to 2005-2012 means, but then increased dramatically in late September 2013, prior to wasting onset in October (Hewson et al., 2018).

Elsewhere, there is evidence that wasting in 2013 – 2014 was tied to elevated primary production. The high pCO_2_ but low temperature-wasting positive relationship noted in Oregon indicates that upwelling may have stimulated primary production at this location (Menge et al., 2016). The CALCoFi program observed highest coastal upwelling on record in 2013 in central California during wasting onset (Leising et al., 2014). These observations suggest that primary production intensity and timing in 2013-2014 departed from inter-annual variation in prior years, and has followed seasonal patterns since 2014. The discontinuous latitudinal emergence of wasting in 2013-2014 and regional apparent longshore sequence of SSW occurrence is consistent with regional and basin-scale patterns of organic matter availability. The spatial scale of phytoplankton blooms sustained solely by terrestrial runoff and groundwater discharge ranges from 880-3600 km^2^ in the Southern California Bight (Santoro et al., 2010). Assuming these blooms are constrained within 10 km of shore, the areal extent of phytoplankton-derived organic matter inputs is well within the reported longshore spread of SSW (Hewson et al., 2014). Upwelling, on the other hand, may affect wider coastal productivity patterns. In 2013, strong upwelling was recorded between 36°N and 48°N (i.e. 1,332 km). It is interesting to note that mass mortality in *Heliaster kubiniji* in the Gulf of California occurred during a prolonged period of heavy rainfall and elevated temperatures prior to El Niño (Dungan et al., 1982). Such rainfall may have caused elevated terrestrial discharge, which in turn may have fueled primary production.

Another potential source of organic matter fueling BLODL is macroalgal detritus (Krumhansl and Scheibling, 2012) and exudates (Abdullah and Fredriksen, 2004). Macroalgae experience seasonal increases in biomass during spring and fall due to elevated temperature and elevated nutrient conditions, but may also experience nutrient limitation in summer (Brown et al., 1997). For example, a study by (Van Alstyne, 2016) found that *Ulva lactuca* abundance was greatest in July when compared to both May and September in Penn Cove. Interestingly the author found that the primary source of nutrients for algal growth during the summer and fall was water from a nearby river, as well as wastewater effluent from a facility at Coupeville (Van Alstyne, 2016). *Laminaria hyperborean* in the North Atlantic Ocean has a pronounced seasonal productivity cycle including an active growing season from February through May during which previous year lamina are shed, followed by a non-growing season until November (Kain, 1979). Exudation of dissolved OM is greatest during *L. hyperborean*’s non-growing season (Abdullah and Fredriksen, 2004). Both macroalgal detritus (Robinson et al., 1982) and exuded OM (Zhang and Wang, 2017) are highly labile and rapidly assimilated by bacteria. While there have been no studies of seasonal exudation or detrital release in the regions affected by SSW, the seasonal variation of SSW and coherence with OM production and detrital breakdown (Krumhansl and Scheibling, 2012) warrants further investigation.

OM originating from decaying asteroids may also generate BLODL. Experimental challenge with asteroid tissue homogenates in this study (Fig. 8), and reported previously (Hewson et al., 2014;Bucci et al., 2017), suggest that wasting may also be associated with decomposition of nearby asteroid individuals via assimilation of tissue-derived compounds and subsequent BLODL. We previously isolated heterotrophic bacteria using sea star tissue homogenates as nutritional source (Hewson et al., 2018). These bacteria include well-known copiotrophic genera. Enrichment of near-benthic OM pools by wasting-affected individuals may have resulted in the apparent density dependence of wasting observed in 2014 in some populations (Hewson et al., 2014). Indeed, challenge with tissue homogenates by direct injection into coelomic cavities likely enriches within-and near animal organic matter pools, which in turn may stimulate heterotrophic remineralization. Hence, challenge experiments, such as those performed previously (Hewson et al., 2014;Bucci et al., 2017) and in this study, may be a consequence of BLODL induced by organic matter availability (and possibly protein-bearing material). The apparent transmissibility of SSW in field sites is based on observations of density dependence at some sites, along with geographic spread between adjacent sites and through public aquaria intake pipes (Hewson et al. 2014). These observations may be inaccurately ascribed to transmissible pathogenic microorganisms, since they may also be explained by enrichment of surrounding habitats and through intake pipes of organic matter pools from decaying individuals.

### Wasted asteroids in 2013-2017 bore stable isotopic signatures of anaerobic processes

Because wasting has no pathognomic signs and has been reported for over a century (reviewed in Hewson et al., 2019), an obvious question is whether BLODL was related to asteroid mass mortality observed from 2013. While retrospective analyses of O_2_ status of asteroids during this event is not possible, hypoxic conditions impart elemental signatures in tissues of preserved specimens. We examined the natural abundance of stable isotopes comparing wasting-affected and grossly normal individuals at the same location and time, in 2013 and 2014. The natural abundance of 15N (δ^15^N) and δ^13^C varied between species, with highest values for *Hippasteria spinosa* and lowest for *Pteraster tesselatus* (Fig. S13). There was no correspondence between known diet of asteroids (and, hence, food web position) and relative stable isotope composition between species. The elemental composition of asteroids, like all animals, largely reflects nutritional source, who obtain anabolic material from consumed prey. Furthermore, asteroids may take up DOM directly from the water column and use these materials for soft body parts, like tube feet (Ferguson, 1967b;a). The half-life of isotopic signatures in tissues relates to tissue turnover and is most stable in ectotherms (Vander Zanden et al., 2015). Dissimilatory anaerobic nitrogen cycling processes, such as denitrification, shift the balance between ^15^N and ^14^N (i.e. selecting against ^15^N), resulting in higher δ^15^N (ratio of tissue ^15^N to atmospheric ^15^N) in environments. Thus, we restricted our analysis of tissue δ^15^N to fast-growing, regenerative tube feet which will therefore reflect the most recent environmental conditions prior to collection. Wasting asteroids (including *Pisaster ochraceus*, *Pycnopodia helianthoides*, and *Evasterias troschelii*), had generally higher δ^15^N in their tissues than asymptomatic tissues at the same site within-species (ns) except for *Leptasterias* sp., which had significantly lower δ^15^N in wasting tissues than in asymptomatic individuals (Fig. 11). On average, δ^15^N was enriched by 3.9 ± 3.3 % for each species (7.0 ± 2.8 % excluding *Leptasterias* sp.) in wasted compared to asymptomatic stars. Ellipse analysis, which can be used to infer isotopic niches or metabolic differences between populations (Jackson et al., 2011) suggested that in all paired site-species comparisons wasted stars have altered C and N metabolisms compared to asymptomatic individuals (Fig. S14).

**Fig. 11:**
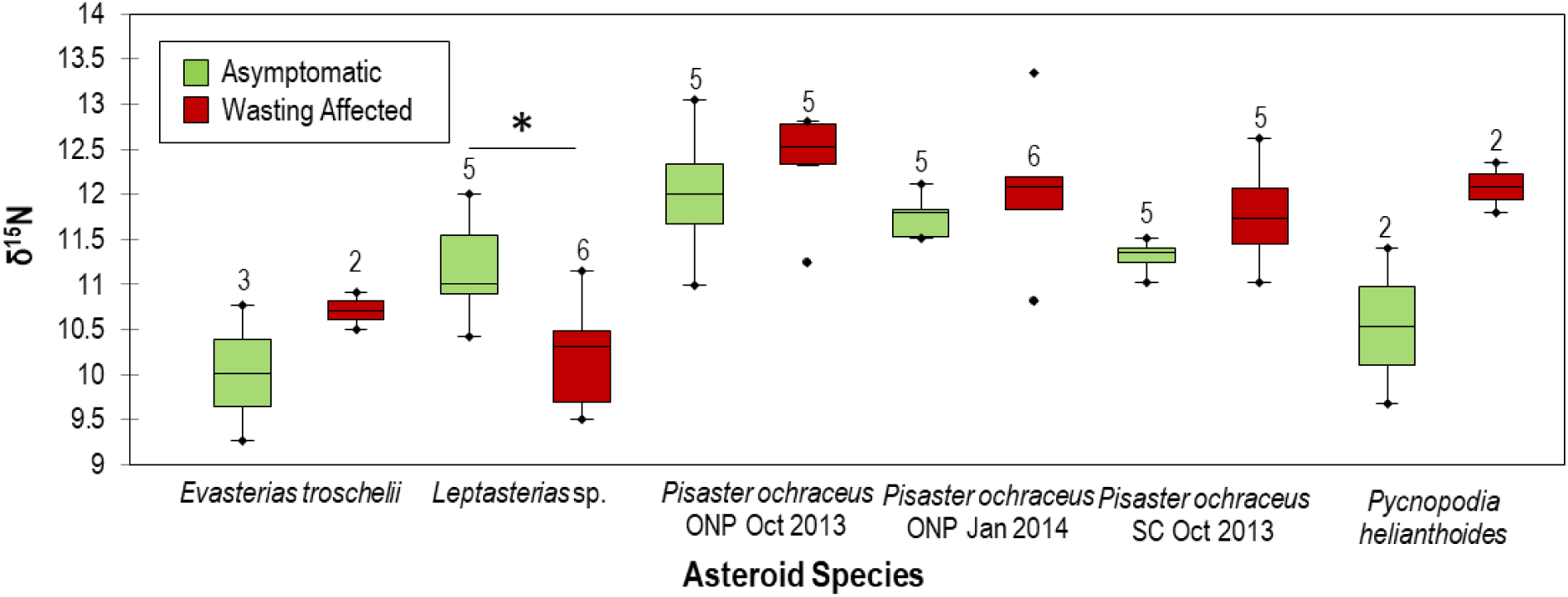
Comparison of asymptomatic and wasting δ^15^N values between species. ONP = Starfish point, Olympic national park and SC = Davenport, Santa Cruz, CA. * indicates p< 0.05. Numbers above box plots indicate *n* of specimens used in comparison.

Translocation of consumed elements to growing tissues is accomplished through continual flux from digestive glands to these tissues through coelomic fluid (Ferguson, 1964). Internal tissues of asteroids are inhabited by a suite of bacteria and archaea (Jackson et al., 2018) including abundant spirochaetes (Holland and Nealson, 1978;Kelly et al., 1995;Kelly and McKenzie, 1995;Nakagawa et al., 2017). Hence, asteroid tube feet tissues, which are distal from digestive glands, may be influenced by heterotrophic microbial activities which enrich for ^15^N over ^14^N. Our finding of higher δ^15^N in most wasted asteroids supports the hypothesis that wasting is associated with enhanced anaerobic dissimilatory respiration of nitrogen species, perhaps during translocation of materials between organs or tissues within asteroids, (Ferguson, 1964) or during uptake of enriched ^15^N in DOM pools surrounding affected asteroids (Ferguson, 1967b;a).

To the best of our knowledge, there has only been one previous report on the effects of hypoxia on stable isotopic composition in animal tissues. Oysters affected by hypoxic water conditions demonstrated δ^15^N enrichment, which they propose was due to hypoxia-induced starvation responses resulting in recycling of internal tissues (Patterson and Carmichael, 2018). Asteroids likewise have similar autophagous responses to starvation, prioritizing somatic maintenance over reproduction (AQUINAS and P., 1976;Harrold and Pearse, 1980). Under typical food availability, reproductive and digestive tissues demonstrate inverse relationships in overall size relating to spawning and feeding time in most asteroid species. However, the ratio between reproductive and digestive tissues in *Leptasterias* spp. is synchronous over time in females (but not so in males), which is different from other starfish species (Menge, 1975). We speculate that the lower δ^15^N observed in wasting *Leptasterias* spp., an opposite trend to other species, may relate to timing of autophagous transfer of materials within individuals and timing of predicted hypoxia (which peaks in late summer) relative to autophagy within animals. It is also possible that asymptomatic and wasting affected specimens were different species of *Leptasterias* sp. since they form a cryptic species complex (Melroy et al., 2017), which may affect comparison between disease states.

### Further evidence for BLODL association with wasting

Wasting imparts transcriptional and population genetic changes in asteroids and surviving populations, respectively. In a metatranscriptomic studies comparing gene expression between wasting and asymptomatic individuals, the relative transcription of high affinity cytochrome c oxidase (ccb3; Preisig et al., 1996) was higher in symptomatic individuals (Gudenkauf and Hewson, 2015). Furthermore, cytochrome P450 2J6, which plays a dual role in both oxidation and detoxification of H_2_S (Tobler et al., 2014), was expressed in at least two studies of wasting asteroids (Fuess et al., 2015;Gudenkauf and Hewson, 2015). Surviving juvenile recruits are genetically distinct to asteroids before 2013 (Schiebelhut et al., 2018). Loci selected for in surviving populations correspond to those heightened in experiments with elevated temperature (Ruiz-Ramos et al., 2020). In particular, Ruiz-Ramos et al (2020) found a synchronous decrease in expression of ND5 (NADH dehydrogenase 5) among field-wasting specimens and those subject to temperature challenge in aquaria, and corresponding mutation in ND5 in surviving populations. Extracellular hypoxia causes downregulation of NADH dehydrogenase in vertebrate cells (Piruat and López-Barneo, 2005), and variation in mt ND5 genes is related to hypoxia sensitivity in humans (Sharma et al., 2019). Elevated temperatures may reduce overall O_2_ concentrations in seawater and cause faster microbial growth rates. Hence, previous observations of enhanced temperature corresponding to wasting (Eisenlord et al., 2016;Kohl et al., 2016;Montecino-Latorre et al., 2016;Miner et al., 2018;Harvell et al., 2019) and with periodic temperature excursion frequency (Aalto et al., 2020) are consistent with the BLODL model proposed in our work.

## Conclusion

Here we present evidence to support our hypothesis that wasting is a sequela of BLODL. We provide evidence that this condition may relate to bacterial abundance/compositional shifts on asteroid respiratory surfaces, and that this results from enrichment with OM. While we cannot definitiely identify a specific source of OM that may lead to BLODL, inter-annual wasting in the field corresponds with phytoplankton biomass, and in controlled experiments we demonstrate that algal-derived OM stimulates wasting. We also provide evidence for this effect as occurring in specimens from the 2013 – 2014 mass mortality event. BLODL may be exacerbated under warmer ocean conditions, or conditions in which labile OM from terrestrial sources (which may include anthropogenic nutrient pollution) may be present in coastal environments. Holothurian wasting, bearing similarity to asteroid wasting in gross disease signs, was reported in the Puget Sound and southeast Alaska beginning in 2014 concomitant with asteroid mass mortality (Hewson et al., 2020) suggesting that this phenomena may affect sympatric benthic invertebrates. Most urchin diseases are associated with diverse bacteria capable of anaerobic metabolism (reviewed in Hewson, 2019). Hence, BLODL may help explain the variation in etiologies observed between echinoderms and between other invertebrate groups, especially those that rely on diffusion for respiratory activities.

## ACKNOWLEDGEMENTS

The authors are grateful to Jim Nagel (Penn Cove Shellfish Company), Karl Menard (Bodega Bay Marine Laboratory), Taylor White (UC Santa Cruz), Joe Gaydos, Lizzy Ashley (Seadoc Society), Martin Haulena (Vancouver Aquarium), Lesanna Lahner (Minnesota Zoo) and Kipp Quinby for provision of specimens and data; Betsy Steele (UC Santa Cruz), Joel Markis and Marnie Chapman (University of Alaska Southeast) for use of laboratory space; Kim Sparks and Elliot Jackson (Cornell University) for laboratory assistance; Mary Sewell (U Auckland), Christopher Mah (Smithsonian); Tom Mumford (University of Washington) for helpful conversations on macroalgae; and Thierry Work (USGS) for assistance with experiments and comments on an early manuscript draft. Asteroid collections were performed under permits CF-19-107 from the Alaska Department of Fish and Game, 19-149a from the Washington State Department of Fish and Wildlife and SC-13144 from the California Department of Fish and Wildlife. This manuscript has been released as a pre-print at biorxiv (Aquino et al., 2020). This work was supported by US National Science Foundation Grants OCE-1537111 and OCE-1737127 awarded to IH and USGS Contract G19AC00434 awarded to IH and T. Work. The authors declare no conflict of interest in publication of this work.

## Author Contributions

LMS and IH designed research; CA, RMB, CMD, JK, IRP, PR, JER, LMS, JPS and IH performed research; CA, RMB, CMD, IRP, PR, JER, LMS, JPS, JPW and IH conducted review and wrote the manuscript; PR and IH provided funding for the research.

