## Supplemental Figures and Tables for "Evidence that non-pathogenic microorganisms drive sea star wasting disease through boundary layer oxygen diffusion limitation"

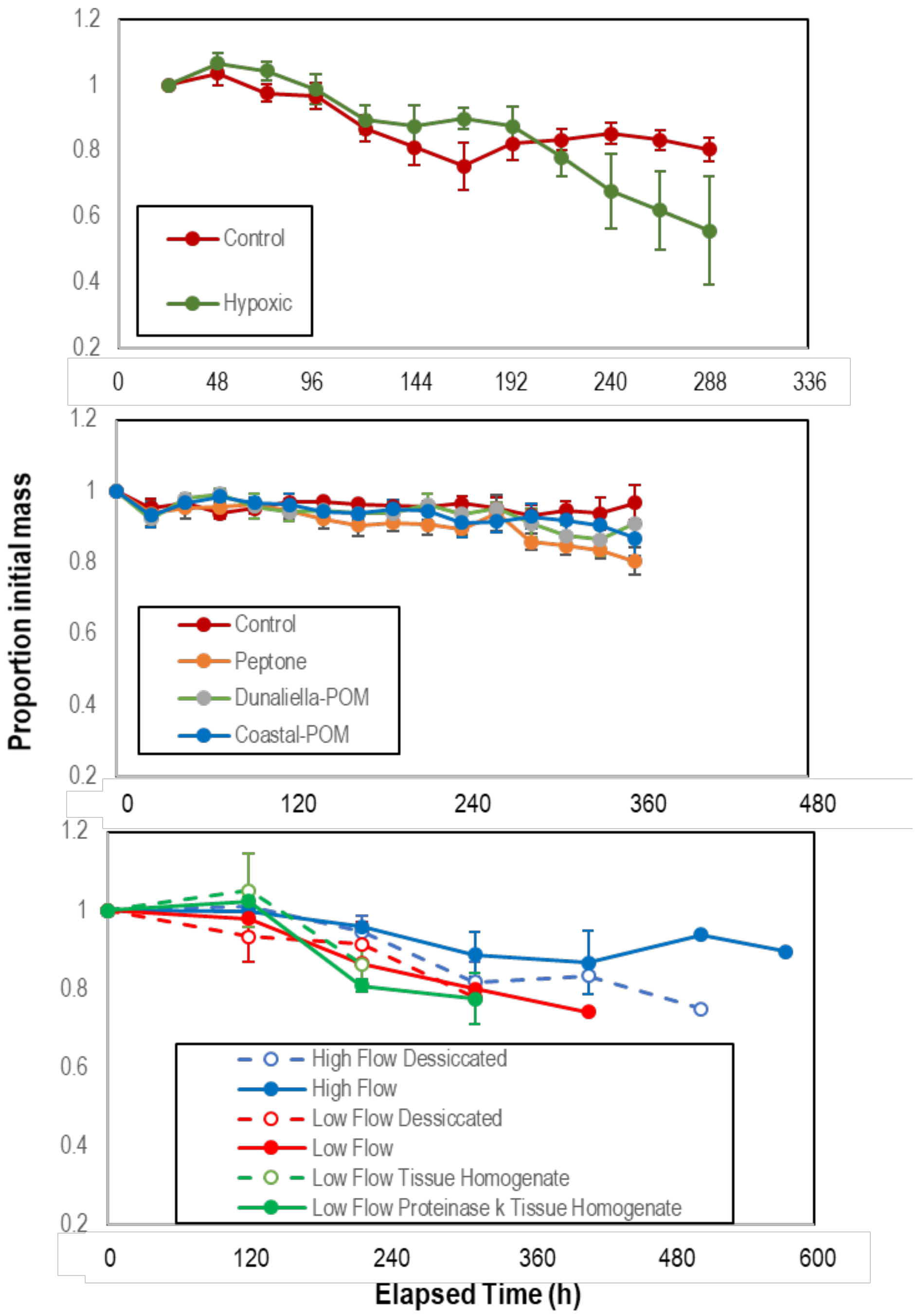


**Fig. S1:** Proportion original mass of individuals exposed to treatments with hypoxia (top), organic substrates (middle) and flow rates/desiccation and tissue homogenates (bottom) during challenge experiments. There was no significant difference in the rate of mass loss in any treatment within each experiment.

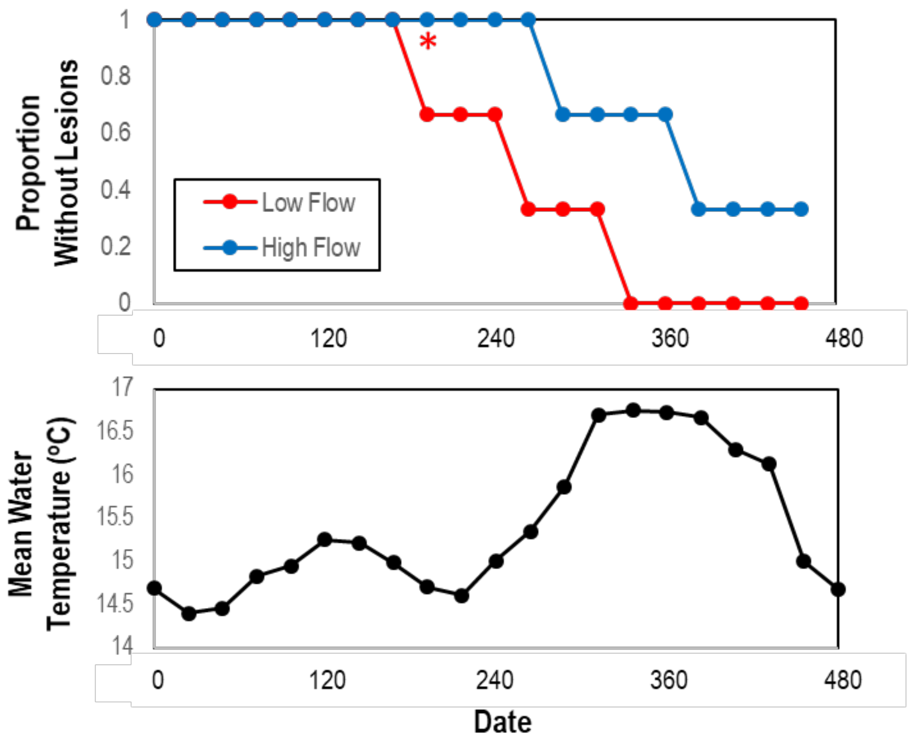


**Fig. S2:** Proportion asymptomatic of *P. ochraceus* (n = 3) and mean water temperature over time subjected to high or low aquarium water flow rates. * denoted that the time of lesion formation was significantly shorter in the low flow rate incubations than in high-flow incubations (Student's *t*-test).

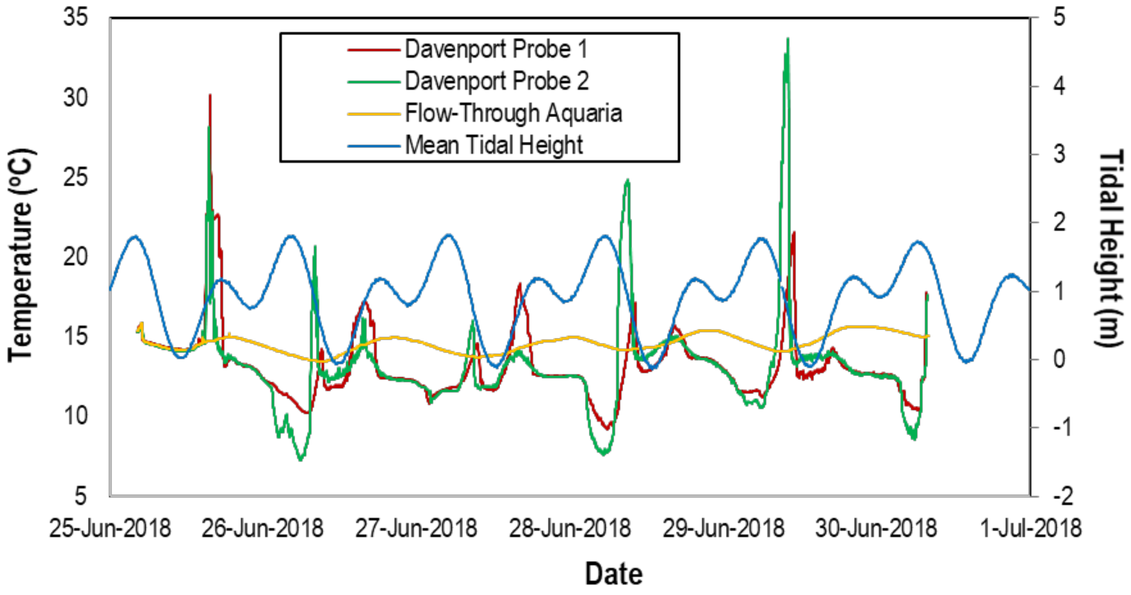


**Fig S3:** Temperature of habitat in which intertidal sea stars exist at Davenport, CA, as measured by Onset HOBO continuous oxygen meters and in flow-through aquaria at the Long Marine Laboratory. Tidal height is also indicated.

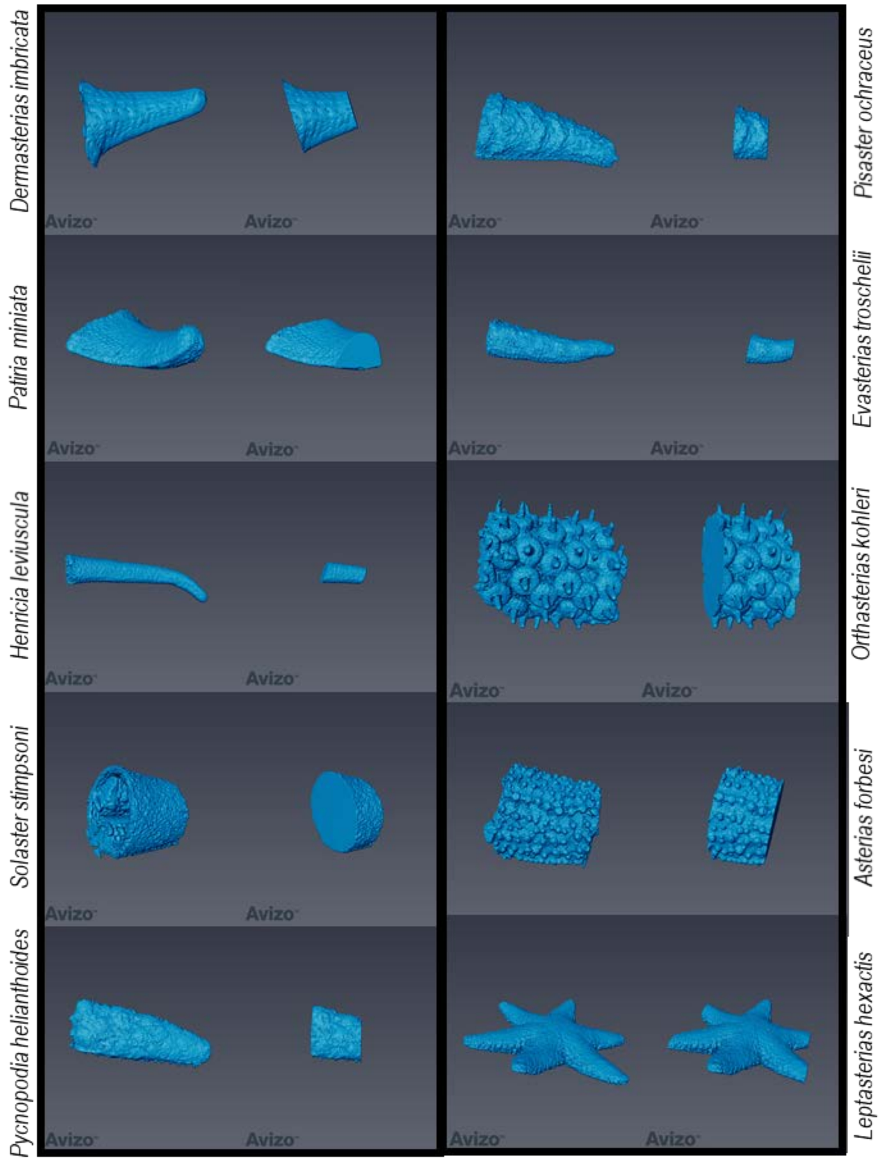


**Fig. S4:** Micro-computed tomographic images of asteroid rays used to examine rugosity. For each species, the entire ray (left) and the 1cm slice of the ray (right) is depicted.

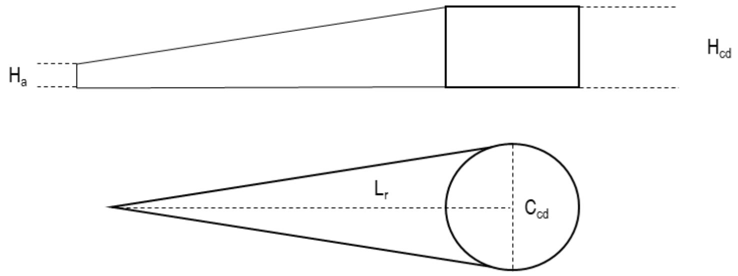


Central disc area ($A_{cd}$) = $\pi(C_{cd}/2)^2$

Height-adjusted ray length ($L_{radj}$) = $\sqrt{(H_{cd}-H_a)^2+(L_r-[C_{cd}/2])^2}$

Total number of rays (a)

Ray area ($A_a$) = $L_{radj}\,[2\pi(C_{cd}/2) / a\,] / 2$

Total 2 dimensional area $A_{2D} = A_{cd} + aA_a$

**Fig. S5:** Calculation of two dimensional area from parameters measured on starfish. Three dimensional area was determined by computed tomography.

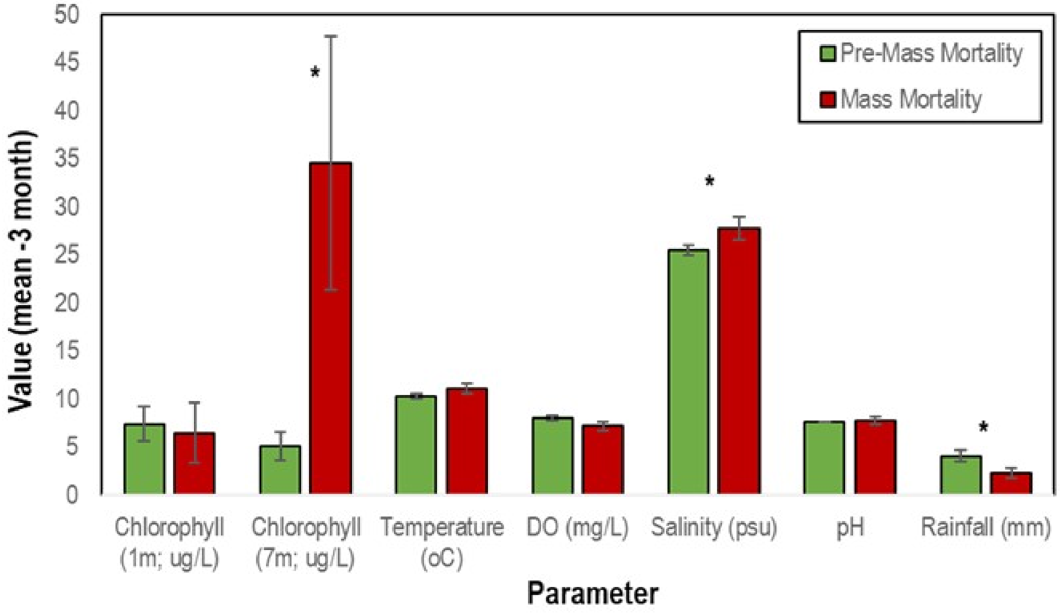


**Fig. S6:** Comparison of physicochemical, meteorological parameters and chlorophyll a concentration at 1 and 7m depth in the 3 months prior to and including the month of mass mortality with months that did not experience mass mortality. * indicates $p<0.05$.

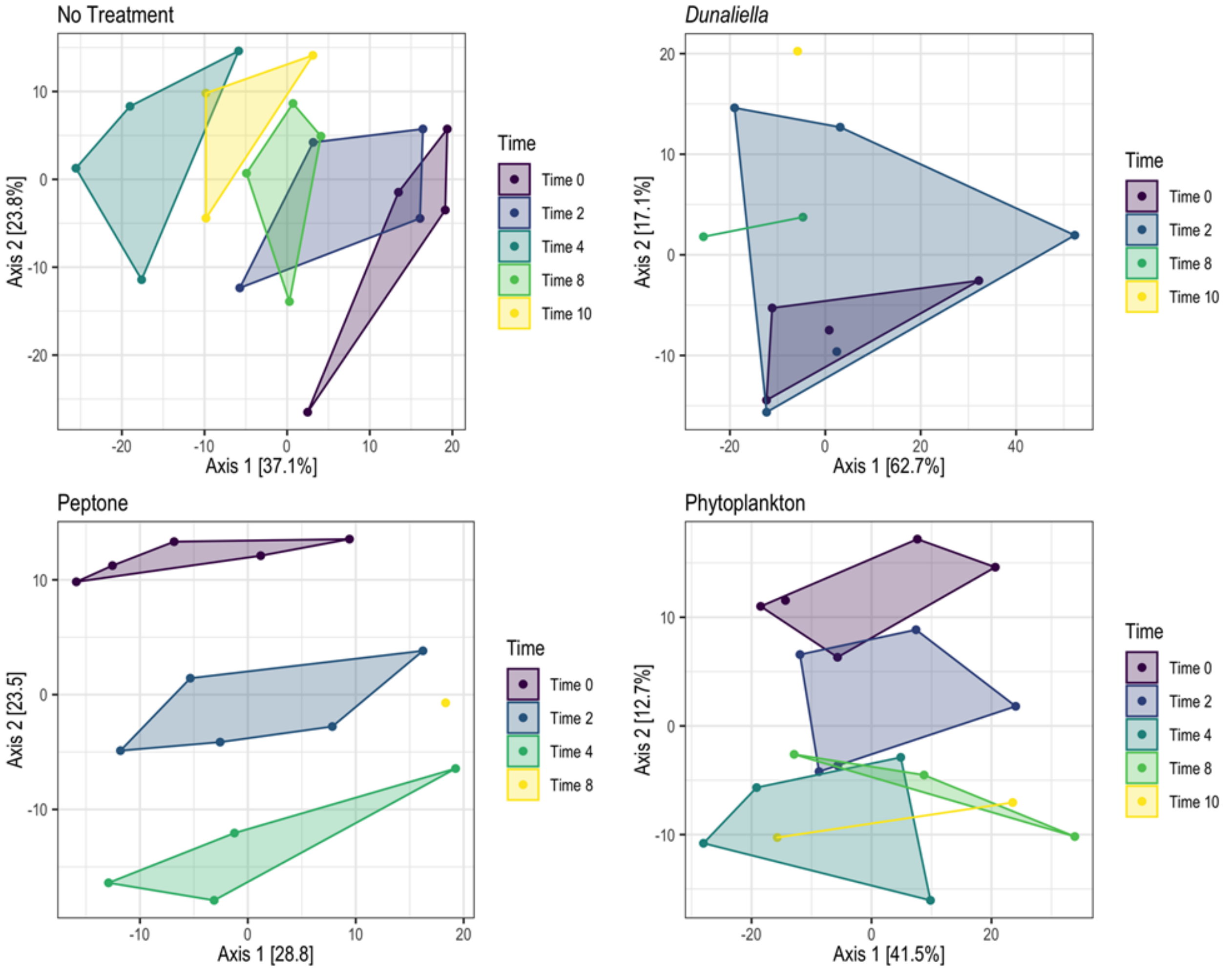


**Fig. S7:** Principal coordinate analyses (PCoA) of Euclidean distances after PhILR transformation. Bacterial communities were derived from surface swabs of *P. ochraceus* specimens enriched with the indicated organic material. Samples were taken over a timecourse indicated in the PCoA legends. Time 0 = initial samples; Time 2 = 48 h, Time 4 = 96 h, Time 8 = 192 h and Time 10 = 240 h. Organic amendment revealed much more variation before lesions first appeared than in body wall or epidermal samples taken at the time of lesion formation or afterwards.

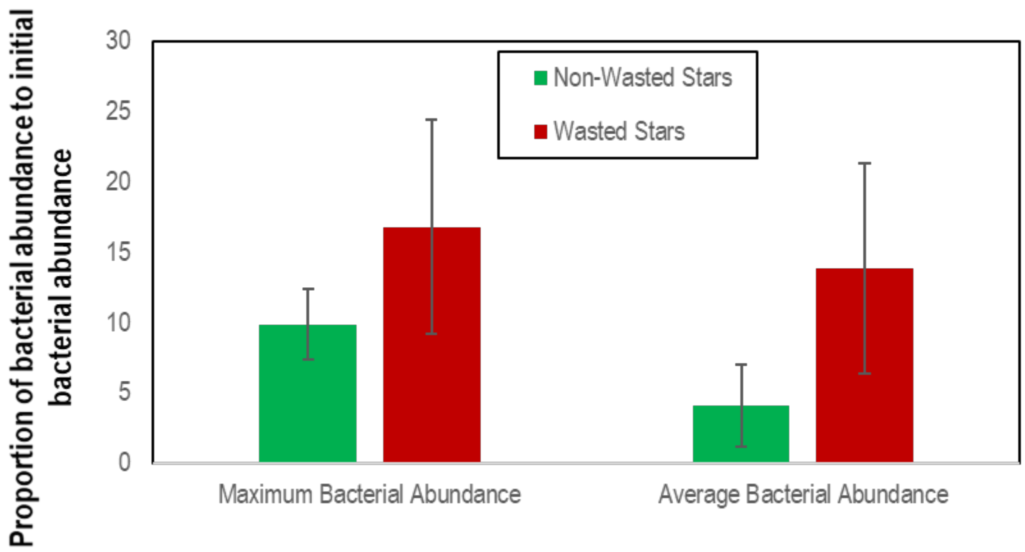


**Fig. S8:** Mean (±SE) proportion of maximum and average bacterial abundance to initial abundance on non-wasted stars and wasting-affected stars (regardless of treatment) during organic matter amendment experiment.

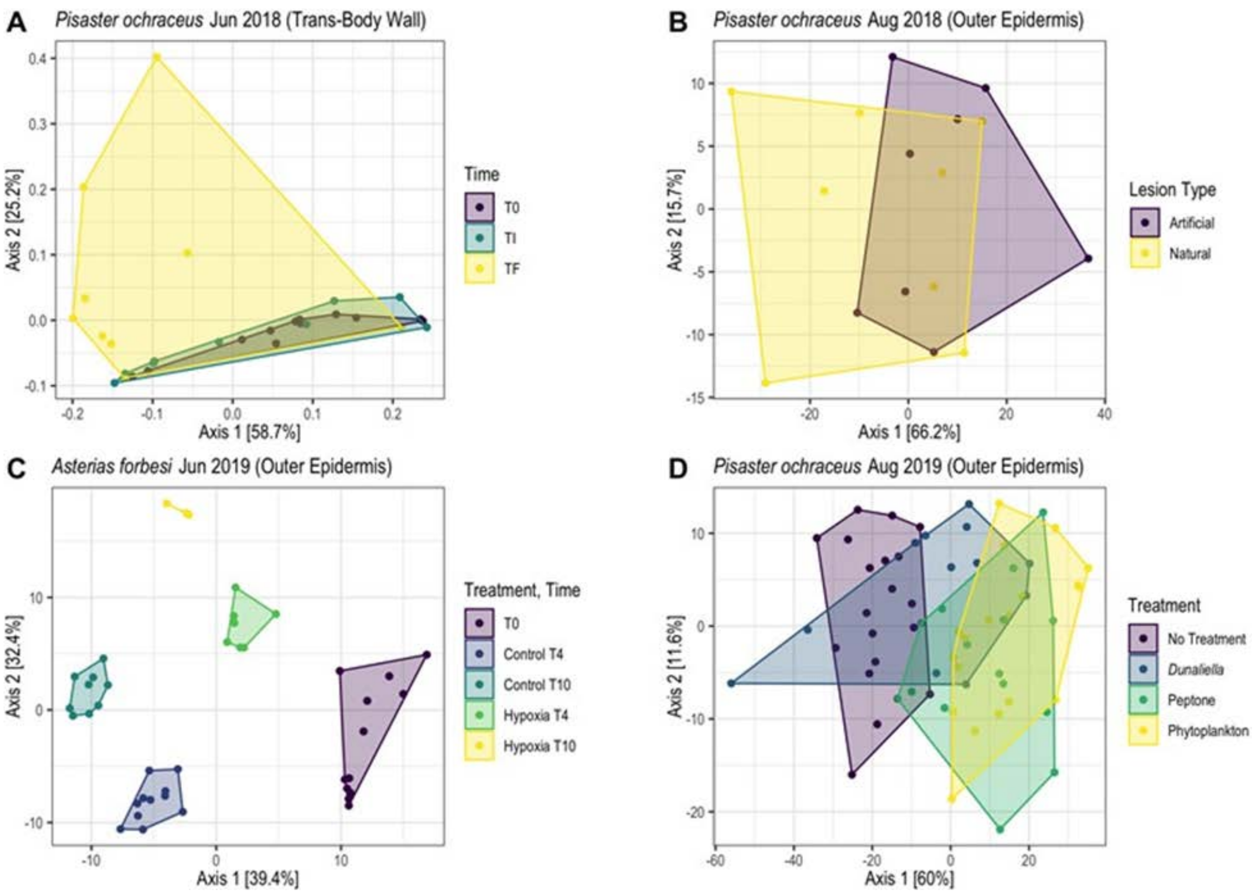


**Fig S9:** Principal coordinate analyses (PCoA) of bacterial communities using Weighted Unifrac distances (A) and Euclidean distances after PhILR transformation (B-C). (A) Body wall samples over the course of wasting. T0 = experiment commencement; TI = lesion genesis; TF = time of death. (B) Epidermal scrapes from artificially induced and naturally occurring lesions. (C) Surface swabs from asteroids under normoxic (control) and suboxic conditions. T0 = experiment commencement; encompasses both normoxic and suboxic conditions. T4, T10 = 96 hand 240 h, respectively. (D) Surface swabs from asteroids amended with different types of organic matter. The timecourse delineation for each treatment is shown in Fig. S8. In the May-June 2018 experiment, microbial assemblages were different between initial samples, after lesions had formed, and at the time of death in body wall samples (Principal Coordinate Analysis of Weighted Unifrac distances; PERMANOVA; P = 0.002).

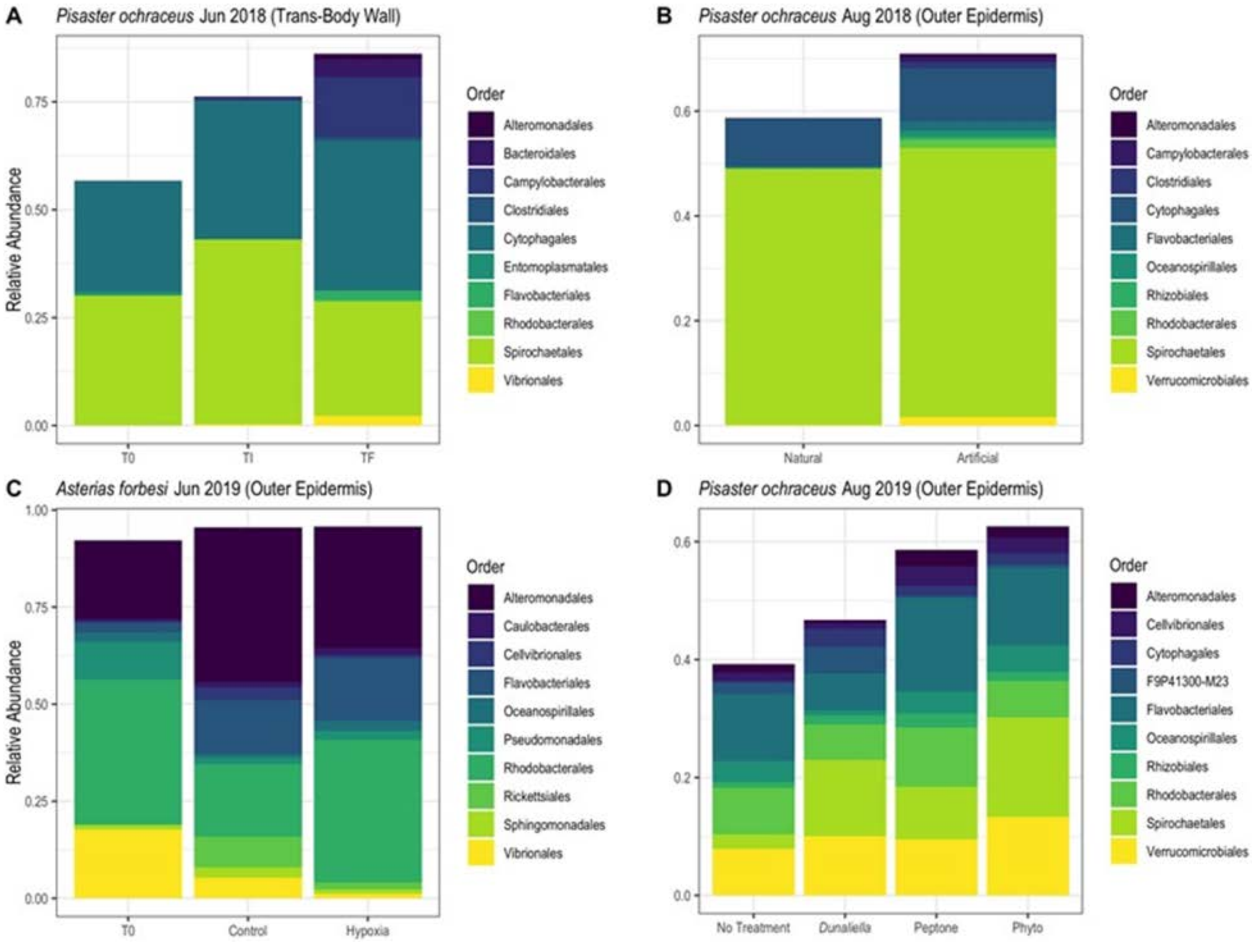


**Fig S10:** Relative abundances of bacterial orders across experiments. (A) Body wall samples over the course of wasting. T0 = experiment commencement; TI = lesion genesis; TF = time of death. (B) Epidermal scrapes from artificially induced and naturally occurring lesions. (C) Surface swabs from asteroids under normoxic (control) and suboxic conditions. T0 represents all asteroids in both conditions at the beginning of the experiment. (D) Surface swabs from asteroids amended with different types of organic matter. Each bar in (C) and (D) (with the exception of T0 in (C)) is an amalgamation of all respective samples collected over the timecourse. The relative abundance of taxa within asteroid-associated microbiomes varied with sample type. Body wall (i.e. using a biopsy punch; May-June 2018) and epidermis of lesion margin (i.e. scrape on margin; August 2018) assemblages were dominated by orders Spirochaetales and Cytophagales, while assemblages immediately above and on surfaces (collected by swabs), bore a large proportion of orders associated with organic rich environments, e.g., Alteromonadales, Flavobacteriales, Rhizobiales, and Rhodobacterales.

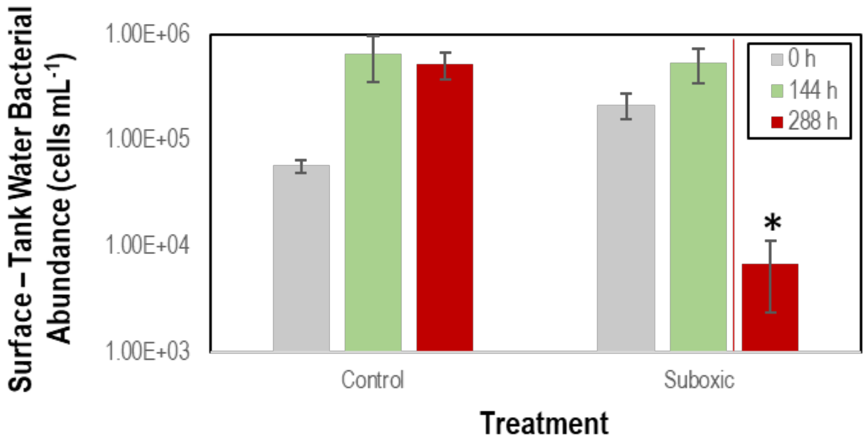


**Fig. S11:** Bacterial abundance in surface layer above *Asterias forbesi* during hypoxic and control treatments (n = 12 each). The abundance was corrected for aquarium water bacterial abundance and measured by SYBR Gold epifluorescence microscopy. The solid vertical red line indicates the average time that suboxic-treated asteroids formed lesions. * denotes $p<0.05$ (Student's *t*-test)

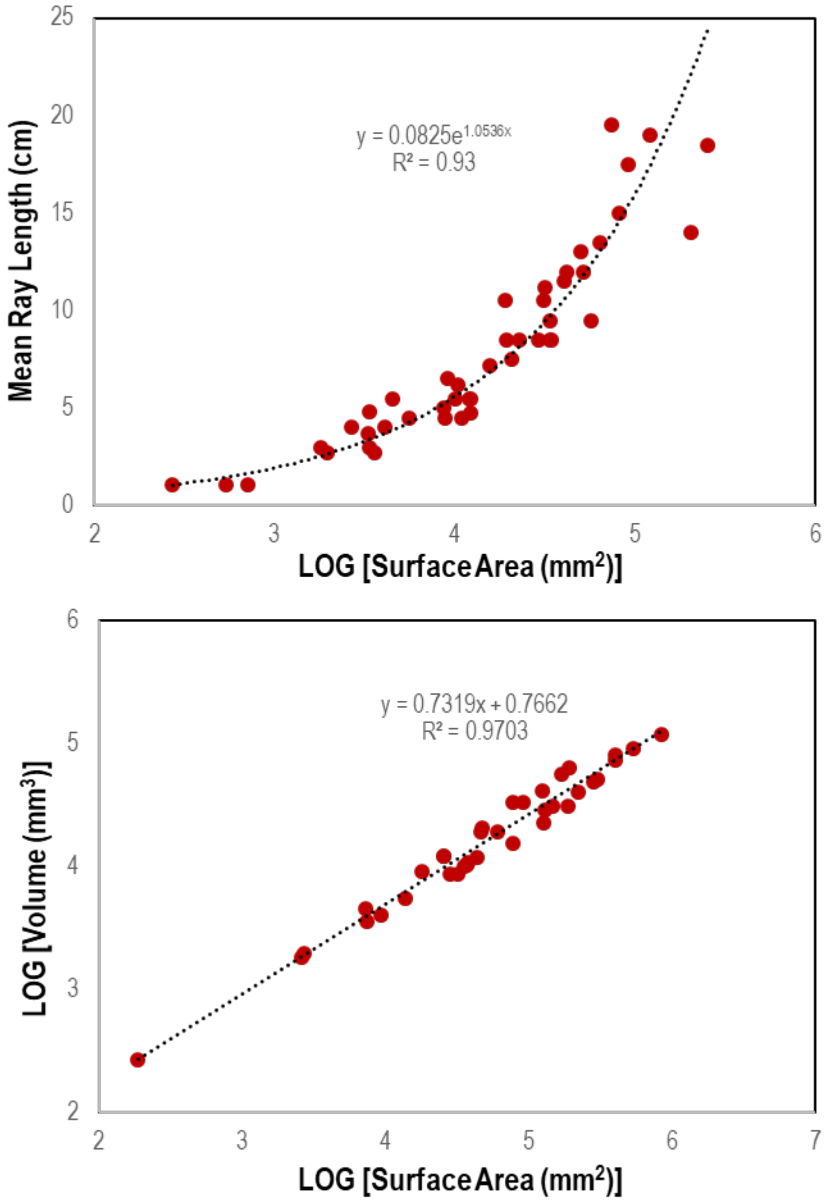


**Fig. S12**: Relationship between ray length and total surface area, and between volume and surface area of 36 specimens of asteroid.

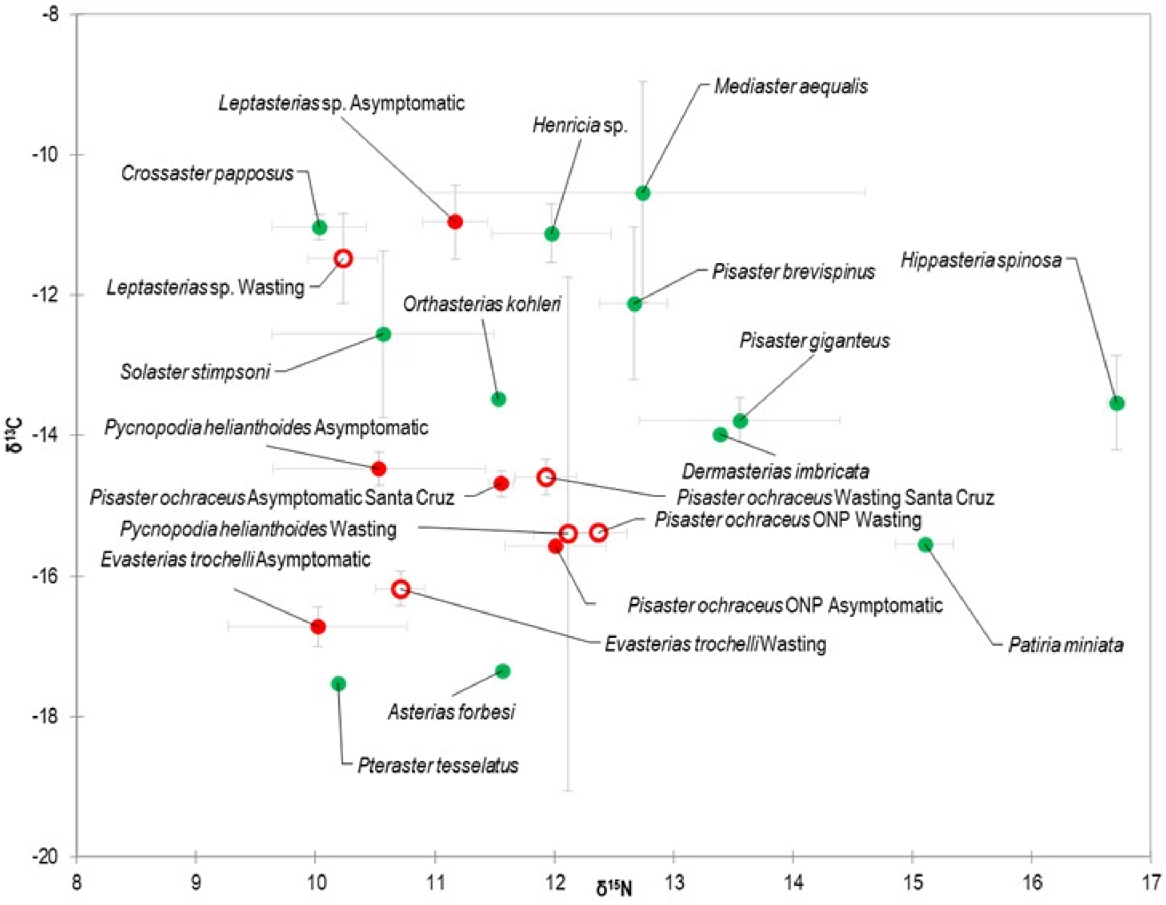


**Fig. S13:** Stable isotopic ($\delta^{13}$C and $\delta^{15}$N) composition of asteroids affected by SSW compared to those less affected by wasting. Green data points indicate species for which only asymptomatic tissue samples were available, while red symbols indicate samples for which both asymptomatic (filled) and wasting-affected (open) tissue samples were analyzed.

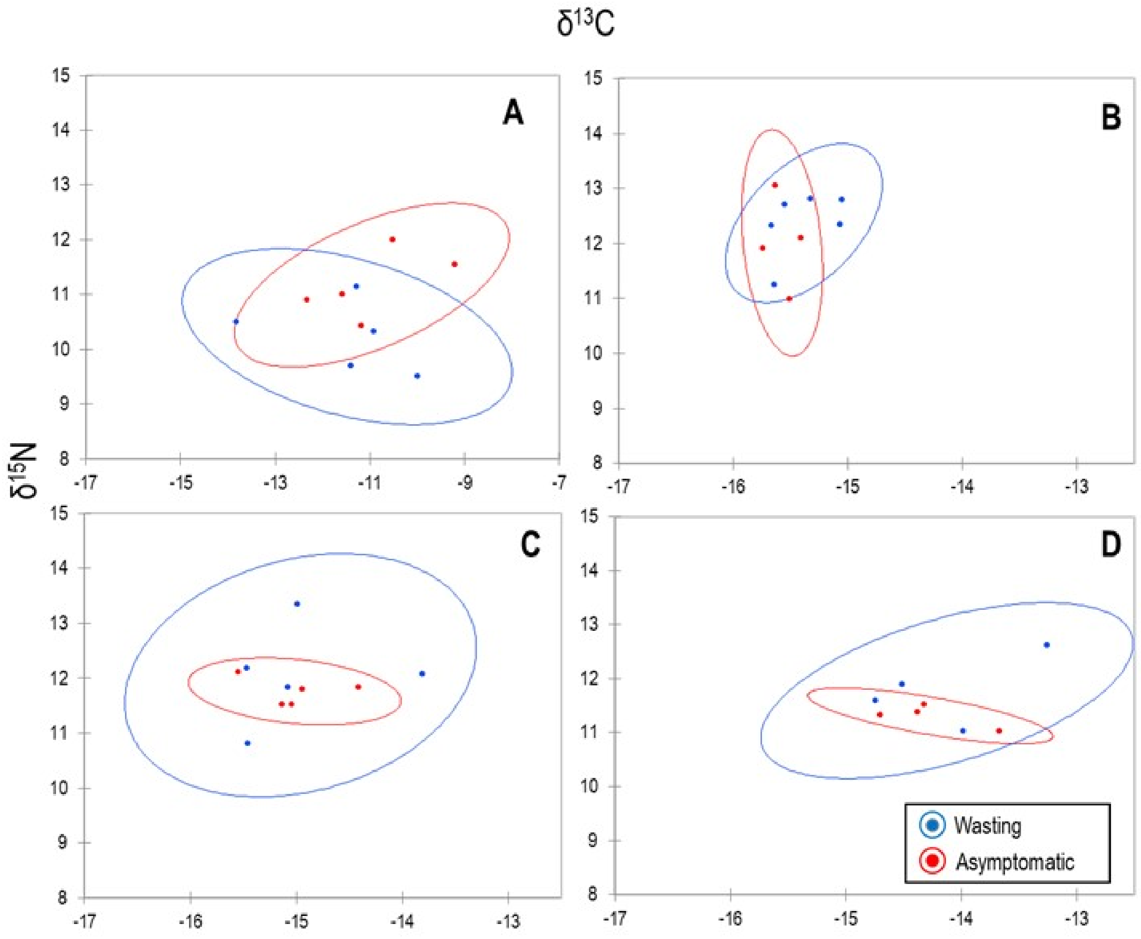


**Fig. S14:** Comparison of $\delta^{13}C/\delta^{15}N$ between asymptomatic and wasting-affected asteroids. Ellipses indicate 95% confidence interval (chi-squared distance). A: *Leptasterias hexactis*, Sitka January 2014; B: *Pisaster ochraceus* Olympic National Park Oct 2013; C: *Pisaster ochraceus* Santa Cruz Oct 2013; D: *Pisaster ochraceus* Olympic National Park Jan 2014

**Table S1:** Metadata on animal size, mass, aquarium flow rate, and change in mass over time for high/low flow, desiccation and tissue homogenate experiments performed in Santa Cruz, CA, June 2018.

| Flow Regime | Treatment | Mass (g) | Ray Length (cm) | Flow Rates (ml s-1) | % Change mass d-1 over experiment | % Change Mass d-1 in First 5 d |
|---|---|---|---|---|---|---|
| High | None | 140 | 8.5 | 6.697924 | -1.681 | 0.4 |
| High | None | 426 | 13 | 6.872852 | -6.72776 | -2.4 |
| High | None | 192 | 10 | 7.267442 | -0.82517 | 0.4 |
| Low | None | 420 | 13 | 2.824859 | -7.37466 | 2.8 |
| Low | None | 186 | 9.5 | 3.532321 | -3.18778 | 0 |
| Low | None | 378 | 13.5 | 3.078818 | -5.84906 | -7.2 |
| High | Desiccation | 464 | 13.5 | 5.889282 | -6.95418 | 2.8 |
| High | Desiccation | 192 | 10 | 8.481764 | -3.82749 | -0.4 |
| High | Desiccation | 192 | 11 | 6.906077 | -2.39557 | 0.4 |
| Low | Desiccation | 492 | 14.5 | 1.835536 | -8.2459 | -19.2 |
| Low | Desiccation | 330 | 13.5 | 3.281917 | 0.8 | 0.8 |
| Low | Desiccation | 642 | 14.5 | 3.313453 | -9.77898 | -2.4 |
| Low | Homogenized Tissue | 438 | 14 | 3.443526 | -8.06557 | 6 |
| Low | Homogenized Tissue | 554 | 16.5 | 3.838772 | -4.62295 | 22.4 |
| Low | Homogenized Tissue | 218 | 10 | 3.371544 | -5.2 | -5.2 |
| Low | Proteinase-k Treated Tissue Homogenate | 498 | 13 | 4.496403 | -9.14754 | 0.4 |
| Low | Proteinase-k Treated Tissue Homogenate | 476 | 11.5 | 3.846154 | -7.50943 | 3.2 |
| Low | Proteinase-k Treated Tissue Homogenate | 180 | 9.5 | 3.772161 | -4.5876 | 1.2 |

**Table S2:** Model parameters used to study interaction between measured variables and lesion time.

| **Model Parameters** | **Parameter Description** | ***P. ochraceus* - Initial Wasting** | ***P. ochraceus* - Flow Rate** | ***P. ochraceus* - OM** | ***A. forbesi*- Oxygen** |
|---|---|---|---|---|---|
| Lesion Time | Time from initiation of experiment to first appearance of lesion | X | X | X | X |
| Mass | Mass of animal | X | X | X | X |
| Ray Length | Average length of ray | X | X | | |
| Aquarium Flow Rate | Flow rate of seawater into incubation container | | X | | |
| Overall Mass Loss | Mass lost over the period of the experiment | | X | X | X |
| 3d Mass Loss | Mass lost over first 3 days of incubation | | | X | X |
| 5d Mass Loss | Mass lost over the first 5 days of incubation | | X | | |
| Surface Area | Surface area of individual (measured by computed tomography) | X | X | X | X |
| Surface Area:Volume | Surface area : Volume of individual (measured by computed tomography) | X | X | | X |
| Respiration Rate | Respiration rate of individual measured at experiment initiation | | | X | X |
| Measured:Hypothetical Diffusion | Respiration rate as a ratio of calculated hypothetical diffusion rate | | | X | X |
| Initial Bacterial Abundance | Bacterial abundance at experiment initiation | | | X | X |
| 3d Bacterial Abundance | Bacterial abundance change in first 3 d of experiment | | | X | |
| Maximum Bacterial Abundance | Maximum bacterial abundance during experiments | | | X | X |

**Table S3:** Species and metadata for intact whole animal specimens collected for computed tomography.

| Species | Date of Collection | Location of Collection | Latitude (ºN) | Longitude (ºW) | Ray length of Individual (cm) |
|---|---|---|---|---|---|
| *Asterias forbesi* | 5/29/2019 | Bar Harbor, ME | 44.427558 | 68.199568 | 7.5 |
| *Dermasterias imbricata* | 9/4/2019 | Sitka, AK | 57.076134 | 135.41194 | 3 |
| *Dermasterias imbricata* | 9/4/2019 | Sitka, AK | 57.076134 | 135.41194 | 8.5 |
| *Dermasterias imbricata* | 9/4/2019 | Sitka, AK | 57.076134 | 135.41194 | 5 |
| *Dermasterias imbricata* | 9/4/2019 | Sitka, AK | 57.098892 | 135.40156 | 8.5 |
| *Dermasterias imbricata* | 9/4/2019 | Sitka, AK | 57.098892 | 135.40156 | 6 |
| *Dermasterias imbricata* | 9/4/2019 | Sitka, AK | 57.098892 | 135.40156 | 4.5 |
| *Dermasterias imbricata* | 9/4/2019 | Sitka, AK | 57.076134 | 135.41194 | 5.5 |
| *Dermasterias imbricata* | 9/4/2019 | Sitka, AK | 57.076134 | 135.41194 | 11.25 |
| *Dermasterias imbricata* | 9/4/2019 | Sitka, AK | 57.076134 | 135.41194 | 7.25 |
| *Evasterias troschelii* | 9/4/2019 | Sitka, AK | 57.076134 | 135.41194 | 3.5 |
| *Evasterias troschelii* | 10/1/2019 | Friday Harbor, WA | 48.595637 | 122.98137 | 19 |
| *Evasterias troschelii* | 9/4/2019 | Sitka, AK | 57.076134 | 135.41194 | 11 |
| *Evasterias troschelii* | 9/4/2019 | Sitka, AK | 57.076134 | 135.41194 | 3 |
| *Evasterias troschelii* | 9/4/2019 | Sitka, AK | 57.076134 | 135.41194 | 10.5 |
| *Evasterias troschelii* | 9/4/2019 | Sitka, AK | 57.076134 | 135.41194 | 2 |
| *Evasterias troschelii* | 10/8/2019 | Friday Harbor, WA | 48.41732 | 122.66429 | 19.5 |
| *Henricia leviuscula* | 9/4/2019 | Sitka, AK | 57.076134 | 135.41194 | 4 |
| *Henricia leviuscula* | 9/4/2019 | Sitka, AK | 57.076134 | 135.41194 | 4.5 |
| *Henricia leviuscula* | 9/4/2019 | Sitka, AK | 57.076134 | 135.41194 | 5 |
| *Henricia leviuscula* | 9/4/2019 | Sitka, AK | 57.076134 | 135.41194 | 3.75 |
| *Leptasterias sp.* | 9/21/2019 | Santa Cruz, CA | 36.626212 | 121.91517 | 1 |

| | | | | | |
|---|---|---|---|---|---|
| *Leptasterias sp.* | 9/21/2019 | Santa Cruz, CA | 36.626212 | 121.91517 | 1 |
| *Leptasterias sp.* | 9/21/2019 | Santa Cruz, CA | 36.626212 | 121.91517 | 0.5 |
| *Orthasterias kohleri* | 9/4/2019 | Sitka, AK | 57.076134 | 135.41194 | 6 |
| *Orthasterias kohleri* | 9/4/2019 | Sitka, AK | 57.076134 | 135.41194 | 8.5 |
| *Orthasterias kohleri* | 9/4/2019 | Sitka, AK | 57.076134 | 135.41194 | 9.5 |
| *Patiria miniata* | 9/21/2019 | Santa Cruz, CA | 36.626212 | 121.91517 | 4.25 |
| *Patiria miniata* | 9/21/2019 | Santa Cruz, CA | 36.626212 | 121.91517 | 4 |
| *Patiria miniata* | 9/21/2019 | Santa Cruz, CA | 36.626212 | 121.91517 | 4.5 |
| *Patiria miniata* | 9/21/2019 | Santa Cruz, CA | 36.626212 | 121.91517 | 2.75 |
| *Pisaster brevispinus* | 10/8/2019 | Friday Harbor, WA | 48.41732 | 122.66429 | 13.5 |
| *Pisaster brevispinus* | 5/24/2016 | Whidbey Island, WA | 48.038582 | 122.40399 | 5.5 |
| *Pisaster giganetus* | 9/21/2019 | Santa Cruz, CA | 36.626212 | 121.91517 | 17.5 |
| *Pisaster giganteus* | 9/21/2019 | Santa Cruz, CA | 36.626212 | 121.91517 | 15 |
| *Pisaster giganteus* | 9/21/2019 | Santa Cruz, CA | 36.626212 | 121.91517 | 23 |
| *Pisaster ochraceus* | 9/4/2019 | Sitka, AK | 57.098892 | 135.40156 | 10.75 |
| *Pisaster ochraceus* | 9/4/2019 | Sitka, AK | 57.098892 | 135.40156 | 10 |
| *Pisaster ochraceus* | 9/4/2019 | Sitka, AK | 57.098892 | 135.40156 | 8 |
| *Pisaster ochraceus* | 9/4/2019 | Sitka, AK | 57.098892 | 135.40156 | 11 |
| *Pisaster ochraceus* | 9/4/2019 | Sitka, AK | 57.098892 | 135.40156 | 13 |
| *Pycnopodia helianthoides* | 9/4/2019 | Sitka, AK | 57.076134 | 135.41194 | 4.5 |
| *Pycnopodia helianthoides* | 9/4/2019 | Sitka, AK | 57.076134 | 135.41194 | 15 |
| *Pycnopodia helianthoides* | 9/4/2019 | Sitka, AK | 57.076134 | 135.41194 | 8.5 |
| *Solaster stimpsoni* | 10/1/2019 | Friday Harbor, WA | 48.595637 | 122.98137 | 18.5 |

**Table S4:** Mass-normalized respiration rates, mean ray length and mean animal mass of asteroids, and ratio of measured to maximum hypothetical diffusion rates. Two non-asteroid species are included for comparison. Respiration rates of both *Asterias forbesi* and *Pisaster ochraceus* were higher than for both *Dermasterias imbricata* and *Patiria miniata* ($p<0.0125$; Student's t-test with Bonferroni correction).

| **Species** | **n** | **Respiration Rate (µmol $kg^{-1}$ $h^{-1}$)** | **Ray Length (cm)** | **Mass (g)** | **Measured:Maximum Diffusion Rate** |
|---|---|---|---|---|---|
| *Asterias forbesi* | 24 | 1287.2 ± 147.2 | 7.7 ± 0.5 | 63.4 ± 6.9 | 2.8 ± 0.5 |
| *Pisaster ochraceus* | 20 | 590.3 ± 26.6 | 11.9 ± 0.4 | 303.6 ± 28.5 | 2.7 ± 0.2 |
| *Dermasterias imbricata* | 2 | 126.1 ± 60.1 | 7.7 ± 0.1 | 196.0 ± 10.0 | 0.7 ± 0.3 |
| *Patiria miniata* | 2 | 46.5 ± 37.0 | 7.8 ± 0.1 | 182.0 ± 20.0 | 0.6 ± 0.5 |

**Table S5:** Species and sampling characteristics of individuals used to compare natural abundance of $^{15}N$ and $^{13}C$ between symptomatic and asymptomatic tissue states and between species.

| Species | Sampling Date | Sampling Location | Disease State | # Individuals |
|---|---|---|---|---|
| *Pisaster ochraceus* | 10/23/2013 | Starfish Point, Olympic National Park | Asymptomatic | 5 |
| *Pisaster ochraceus* | 10/23/2013 | Starfish Point, Olympic National Park | Wasting Affected | 5 |
| *Pisaster ochraceus* | 1/3/2014 | Starfish Point, Olympic National Park | Asymptomatic | 5 |
| *Pisaster ochraceus* | 1/3/2014 | Starfish Point, Olympic National Park | Wasting Affected | 6 |
| *Pisaster ochraceus* | 10/8/2013 | Davenport, CA | Asymptomatic | 5 |
| *Pisaster ochraceus* | 10/8/2013 | Davenport, CA | Wasting Affected | 5 |
| *Pycnopodia helianthoides* | 10/17/2013 | Gibbons, BC, Canada | Asymptomatic | 2 |
| *Pycnopodia helianthoides* | 10/16/2013 | Gibbons, BC, Canada | Wasting Affected | 2 |
| *Evasterias trochellii* | 10/17/2013 | Gibbons, BC, Canada | Wasting Affected | 3 |
| *Evasterias trochellii* | 10/17/2013 | Gibbons, BC, Canada | Asymptomatic | 2 |
| *Leptasterias hexactis* | 9/10/2014 | Sitka, AK | Asymptomatic | 5 |
| *Leptasterias hexactis* | 9/10/2014 | Sitka, AK | Wasting Affected | 5 |
| *Solaster stimpsoni* | 1/8/2016 | Blakley Is, WA | Asymptomatic | 2 |
| *Patiria miniata* | 1/23/2014 | Monterey, CA | Asymptomatic | 2 |
| *Crossaster papposus* | 1/10/2016 | Clinton, WA | Asymptomatic | 2 |
| *Dermasterias imbricata* | 1/11/2016 | Clinton, WA | Asymptomatic | 1 |
| *Henricia sp.* | 1/10/2016 | Clinton, WA | Asymptomatic | 2 |
| *Hippasteria spinosa* | 1/6/2016 | West Point, WA | Asymptomatic | 2 |
| *Mediaster aequalis* | 1/7/2016 | Clinton, WA | Asymptomatic | 2 |
| *Orthasterias kohleri* | 1/10/2016 | Port Townsend, WA | Asymptomatic | 1 |
| *Pisaster brevispinus* | 2/14/2014 | Carpinteria, CA | Asymptomatic | 2 |
| *Pisaster giganteus* | 3/1/2014 | Goleta Pier, CA | Asymptomatic | 2 |
| *Pteraster tesselatus* | 1/8/2016 | Friday Harbor, WA | Asymptomatic | 1 |
| *Asterias forbesi* | 10/3/2013 | Bar Harbor, ME | Asymptomatic | 2 |

**Table S6:** Significance of survival (time to lesion development) in asteroids subjected to treatment under high and low flow regimes. Statistical significance assessed by comparing hazard rate by log rank test.

| | High Flow | High Flow - Desiccated | Low Flow | Low Flow - Desiccated | Low Flow - Tissue Homogenate | Low Flow - Proteinase-k treated Tissue Homogenate |
|---|---|---|---|---|---|---|
| **High Flow** | - | ns | ns | 0.0389 | 0.0246 | 0.0224 |
| **High Flow - Desiccated** | | - | ns | 0.0389 | 0.0246 | ns |
| **Low Flow** | | | - | 0.0389 | ns | ns |
| **Low Flow - Desiccated** | | | | - | 0.0389 | 0.0389 |
| **Low Flow - Tissue Homogenate** | | | | | - | ns |
| **Low Flow - Proteinase-k treated Tissue Homogenate** | | | | | | - |

**Table S8:** Multiple linear regression models of lesion time against measured experimental parameters. LT = Lesion Time; MC = Mass change over experiment, IB = Initial bacterial abundance; FR = Flow Rate; AM = Animal Mass at Experiment Initiation; TB = Bacterial abundance change over first three days of experiment. Models were performed using stepwise (forward selection) regression with change in Akaike's AIC as entry criteria.

| **Experiment** | **Variable 1** | **Variable 2** | **Model** | ***p*** |
|---|---|---|---|---|
| Asterias - Oxygen | MC (p = 0.025) | IB (p=0.036) | LT = 33.78-3.99*IB+42.49*MC | 0.012 |
| Pisaster ochraceus - Flow Rate | FR (p=0.025) | - | LT = 19.46-0.01*AM-0.22*FR | 0.004 |
| Pisaster ochraceus - OM | AM (p=0.011) | TB (p=0.031) | LT = 12.86-0.01*AM-151.72*TB | 0.018 |
| Pisaster ochraceus Flow Rate + OM | AM (p=0.0017) | - | LT = 13.98-0.02*AM-0.28*MC | 0.006 |
